# DRMY1 promotes robust morphogenesis by sustaining the translation of cytokinin signaling inhibitor proteins

**DOI:** 10.1101/2023.04.07.536060

**Authors:** Shuyao Kong, Mingyuan Zhu, M. Regina Scarpin, David Pan, Longfei Jia, Ryan E. Martinez, Simon Alamos, Batthula Vijaya Lakshmi Vadde, Hernan G. Garcia, Shu-Bing Qian, Jacob O. Brunkard, Adrienne H. K. Roeder

## Abstract

Robustness is the invariant development of phenotype despite environmental changes and genetic perturbations. In the Arabidopsis flower bud, four sepals robustly initiate and grow to constant size to enclose and protect the inner floral organs. We previously characterized the mutant *development related myb-like1* (*drmy1*), where 3-5 sepals initiate variably and grow to different sizes, compromising their protective function. The molecular mechanism underlying this loss of robustness was unclear. Here, we show that *drmy1* has reduced TARGET OF RAPAMYCIN (TOR) activity, ribosomal content, and translation. Translation reduction decreases the protein level of ARABIDOPSIS RESPONSE REGULATOR7 (ARR7) and ARABIDOPSIS HISTIDINE PHOSPHOTRANSFER PROTEIN 6 (AHP6), two cytokinin signaling inhibitors that are normally rapidly produced before sepal initiation. The resultant upregulation of cytokinin signaling disrupts robust auxin patterning and sepal initiation. Our work shows that the homeostasis of translation, a ubiquitous cellular process, is crucial for the robust spatiotemporal patterning of organogenesis.

## INTRODUCTION

Robustness, or canalization, is the invariant, reproducible development of phenotype, unchanged by environmental fluctuations, genetic perturbations, or gene expression noise^1–4^. Commonly, within an individual, a given number of organs develop at well-defined positions to a robust final size and shape, which is crucial for fitness under stabilizing selection^2^. For example, transplanted eyes, limbs, and kidneys in mammals grow to a mature size similar to their donor, irrespective of the mature size of the same type of organ in the recipient^5–7^. The pairs of wings and halteres in Drosophila develop to robust, precisely coordinated final size and shape, which are required for flight^8–11^. The characteristic cruciform flower in *Brassicaceae* consists of four petals^12^, a trait that can contribute to pollinator attraction^13^. The robust positioning of leaves around the shoot apical meristem in plants, or phyllotaxis, ensures optimal light capture^14–16^. While these examples of developmental robustness have been documented for a long time, the underlying molecular mechanisms have just begun to be unveiled.

Earlier studies looking for genes involved in maintaining robustness have found *HEAT SHOCK PROTEIN 90* (*HSP90*). Mutations of *HSP90* cause a diverse array of phenotypic changes in plants, fruit fly, zebrafish, worm, and humans^3,17,18^. Notably, the display and severity of these changes vary between individuals and even between different parts of the same individual, indicating that developmental robustness is disrupted^17,18^. *HSP90* encodes a protein chaperone which has clients from nearly all developmental and signaling pathways^3^. *HSP90*, therefore, is a hub gene that affects numerous other genes within the gene network^2^. Disruption of such a hub gene would therefore trigger many defects in numerous developmental processes. Similarly, genes involved in central cellular processes such as chromatin remodelling^19–21^, transcription^19,20^, translation^22,23^, and protein degradation^24,25^ are also hub genes, and they have been found to be important for developmental robustness in various systems including fungi, animals, and plants. How these broad-acting hub genes contribute to the robustness of tissue-specific developmental phenotypes is still largely unclear.

We have developed the *Arabidopsis* sepal as a system to elucidate the mechanisms maintaining robustness in organ size and shape^26–28^. Sepals are the outermost floral organs whose function is to enclose buds and protect the developing inner organs, i.e. petals, stamens, and carpels, before the flower blooms. To fulfill this protective function, each flower robustly develops four sepals of equal length, allowing them to close at the top (Figure 1A, top left); these four sepals are of equal width and positioned 90° from each other, leaving no gap on the sides (Figure 1A, middle left). This robustness in sepal size and shape stems from the robust initiation of the four sepal primordia from the floral meristem with precisely coordinated spatiotemporal patterns^27^ (Figure 1A, bottom left). The initiated sepal primordia attain robust final size and shape by spatiotemporal averaging of cellular growth variability during sepal elongation, and synchronous progression of a whole-flower growth termination signal from tip to base^26^. In addition, noise in gene expression must be kept low to ensure sepal size robustness^29^. We previously characterized a mutant in *DEVELOPMENT RELATED MYB-LIKE 1* (*DRMY1*) that develops flowers where the inner organs are exposed due to gaps between sepals^27^. The gaps are caused by variability in sepal development. Specifically, some sepals are shorter than others, leaving gaps on the top (Figure 1A, top right); the arrangement of sepals around the flower deviate from the canonical form such that parts of the flower are not covered by a sepal, leaving gaps on the side (Figure 1A, middle right). This variability in the size, number, and position of the mature sepal originates from the earliest stages of floral development where the spatiotemporal pattern of sepal initiation becomes variable (Figure 1A, bottom right). Variability in sepal initiation, in turn, is driven by the loss of robust patterning of auxin and cytokinin^27^, two plant hormones critical for morphogenesis^30–32^, in the floral meristem before sepal initiation. However, the molecular mechanism through which DRMY1 maintains robust hormone patterning is still unknown.

**Figure 1.**
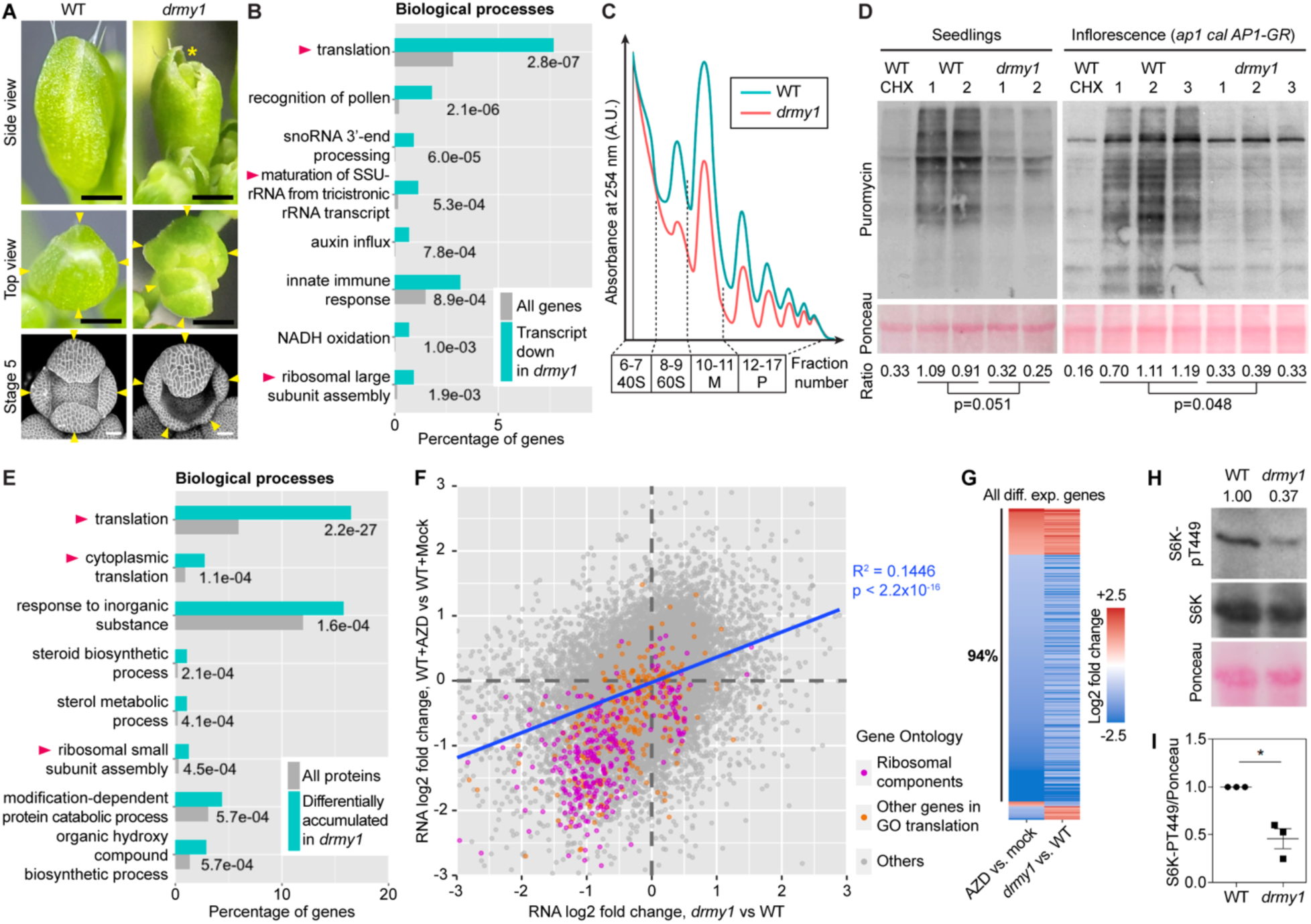
*drmy1* has reduced ribosome abundance, translation rate, and TOR activity. **(A)** Top row, stage 12 buds of WT (left) and *drmy1* (right) viewed from the side. Asterisk shows the gap between sepals with petals and carpels exposed. Middle row, stage 12 buds of WT (left) and *drmy1* (right) viewed from the top. Arrowheads point to sepals. Note that the *drmy1* bud has 5 sepals of unequal size and unevenly spaced, exposing the stamens and carpels. Bottom row, stage 5 buds of WT (left) and *drmy1* (right) containing *35S::mCitrine-RCI2A* (plasma membrane marker). Arrowheads point to sepal primordia. Note that the *drmy1* bud has 5 sepal primordia of different sizes. Scale bars are 0.5 mm for stage 12 bud images and 25 µm for stage 5 bud images. **(B)** Gene ontology (GO) enrichment of downregulated genes in *drmy1* compared to WT, in the *ap1 cal AP1-GR* background. Shown are the top 8 GO terms and their enrichment p-values. A complete list can be found in Supplemental Dataset 1. Arrowheads highlight terms related to ribosome biogenesis or translation. **(C)** Polysomal profiles of WT (blue) and *drmy1* (red) in the *ap1 cal AP1-GR* background, representative of 3 biological replicates each. Additional replicates are in Supplemental Dataset 2. M, monosomes. P, polysomes. **(D)** Puromycin labeling of WT vs *drmy1*. Left, WT and *drmy1* seedlings. From left to right: WT pre-treated with CHX, two biological replicates of WT pre-treated with mock, and two biological replicates of *drmy1* pre-treated with mock. All groups were then treated with puromycin. For seedlings to match in size, WT seedlings were 8 days old and *drmy1* seedlings were 10 days old. Right, WT and *drmy1* inflorescences of induced *ap1 cal AP1-GR* background. From left to right: WT co-treated with puromycin and CHX, three biological replicates of WT treated with puromycin, and three biological replicates of *drmy1* treated with puromycin. In both experiments, RuBisCO large subunit on Ponceau S-stained membrane is shown as a loading control (bottom). Ratio between puromycin and Ponceau S signals, normalized by the mean of WT, is shown on the bottom (p-values are from two-sided Student’s t-test). **(E)** Gene ontology (GO) enrichment of differentially accumulated proteins in *drmy1* compared to WT, in the *ap1 cal AP1-GR* background. Shown are the top 8 GO terms and their enrichment p-values. A complete list can be found in Supplemental Dataset 1. Arrowheads highlight terms related to ribosome biogenesis or translation. **(F-G)** Coherent alteration of gene expression by *drmy1* and AZD-8055 TOR inhibitor treatment. **(F)** Scatterplot of RNA log2 fold change in *drmy1* vs WT (x-axis), and WT+AZD vs WT+Mock (y-axis), in 7-day-old seedlings. Genes are color-coded based on the following categories: genes in “Structural constituents of the ribosome” (GO:0003735) and its offspring terms (magenta); all other genes in “Translation” (GO:0006412) and its offspring terms (orange); all other genes (gray). Blue line shows a linear regression of all points (R^2^ = 0.1446, p < 2.2×10^-16^). **(G)** Of the 466 genes that are differentially expressed under both conditions, 439 (94%) are coherently altered by AZD-8055 treatment and the *drmy1* mutation. **(H-I)** Phosphorylation of the direct TOR substrate, S6K-pT449, in WT and *drmy1*. **(H)** A representative blot. Top, S6K-pT449. Middle, total S6K protein. Bottom, Ponceau S staining. Ratio between S6K-pT449 signal and Ponceau S signal is shown above the blots. **(I)** Ratio between S6K-pT449 and Ponceau S signals normalized by WT, quantified across in three experiments, shows that TOR activity decreased by half in *drmy1*. (mean ± SD; *, p<0.05).

In this study, we elucidate a mechanism through which DRMY1 maintains robust hormone patterning and thus robust sepal initiation. Specifically, we find that DRMY1 maintains proper activity of TARGET OF RAPAMYCIN (TOR), a crucial regulator of ribosome level and mRNA translation^33,34^, and thereby sustains translation *in vivo*. When *DRMY1* is mutated, the levels of ARABIDOPSIS RESPONSE REGULATOR7 (ARR7) and ARABIDOPSIS HISTIDINE PHOSPHOTRANSFER PROTEIN 6 (AHP6), two cytokinin inhibitor proteins that are rapidly synthesized during hormone patterning prior to sepal initiation, are drastically reduced in the floral meristem. Consequently, cytokinin signaling uniformly increases in the meristem periphery, causing variability in auxin patterning and sepal initiation. We further propose that the increase in cytokinin signaling may be a survival mechanism to alleviate the translation rate reduction when ribosomal content is limited. In summary, our work shows that the hub processes of TOR signaling and translation, which occur in every cell, have very tissue-specific roles in maintaining robust organogenesis by sustaining the rapid synthesis of hormone signaling proteins.

## RESULTS

### The *drmy1* mutant has reduced TOR activity, ribosome content, and translation rate

*DRMY1* encodes a MYB/SANT domain protein which may exert transcriptional regulation^27^. To look for differentially expressed genes in *drmy1* which may be candidates underlying variable sepal initiation, we performed RNA-sequencing (RNA-seq) in *drmy1* and wild type (WT) of *apetala1* (*ap1) cauliflower (cal) AP1-GR* background^35,36^. The *ap1 cal AP1-GR* inflorescence produces numerous tightly packed ball-shaped meristems, which, upon induction, synchronously initiate sepal primordia, allowing us to collect large quantities of floral meristems with sepal primordia initiating (Stage 3)^37^ (Figure S1A). We crossed *drmy1* into *ap1 cal AP1-GR* and performed RNA-seq on induced inflorescences of WT and *drmy1* in this background. We detected transcripts from a total of 21,496 genes, of which 1,042 (4.8%) were differentially expressed in *drmy1* (Figure S1B; Supplemental Dataset 1). We found that the 443 genes downregulated at the transcript level in *drmy1* were most enriched in the gene ontology (GO) term “Translation”, a fundamental and ubiquitous cellular process that converts genetic information from transcript to protein. Within this term, genes encoding ribosomal components were most downregulated (Figure S1C). The 443 downregulated genes were also enriched in several other ribosome-related GO terms (Figure 1B). We therefore hypothesized that ribosomal abundance and translation are affected in *drmy1*, potentially altering the accumulation of proteins critical to developmental robustness.

To determine whether and how ribosomal abundance and translation are affected in *drmy1*, we performed polysome profiling in induced inflorescences of WT and *drmy1* in *ap1 cal AP1-GR* background. Compared to WT, all peaks corresponding to 40S, 60S, monosomes, and polysomes are drastically reduced in *drmy1* (Figure 1C; Supplemental Dataset 2). To see whether this reduction in ribosomal content affected *de novo* protein synthesis rate *in vivo*, we performed puromycin labeling. Samples were incubated with puromycin, an amino acid-tRNA analog that is incorporated into nascent polypeptide chains and can be detected using an anti-puromycin antibody to infer global translation rate^38,39^. In both young seedlings and induced *ap1 cal AP1-GR* inflorescences, we found that the puromycin level detected in *drmy1* mutant samples was much reduced compared to WT (Figure 1D), indicating a reduction in global translation rate. We hypothesized that reduced global translation rate should likely result in globally decreased protein levels. For this, we looked at a ubiquitously expressed membrane marker *UBQ10::mCherry-RCI2A*, and found that it had a small (∼25%) but significant decrease in fluorescence intensity in the inflorescence meristem and young floral buds of *drmy1* compared with WT (Figure S1D, E). We also measured its fluorescence intensity in the ribosomal mutant *ul4y* (*rpl4d*)^40^ and we found that the decrease in fluorescence intensity in *drmy1* is even greater than in *ul4y* (Figure S1F, G). Overall, these results show that ribosomal content and translation are indeed reduced in the *drmy1* mutant.

To test how the global repression of translation in *drmy1* impacts its proteome, we extracted total soluble protein from induced inflorescences of WT and *drmy1* in *ap1 cal AP-GR* background and performed mass spectrometry. We identified a total of 5,077 proteins, of which 548 (10.8%) were differentially accumulated in *drmy1* (Figure S1B; Supplemental Dataset 1). These differentially accumulated proteins were enriched in GO terms related to translation and ribosomes (Figure 1E). Despite the overall reduction in ribosomes (Figure 1C), relative to other proteins, ribosomal components are more abundant in *drmy1* (Figure S1H; Supplemental Dataset 1). This is not true for all proteins involved in translation; poly-A binding proteins and tRNA synthetases, for example, are relatively less abundant in *drmy1* than in WT. Moreover, the 26S proteasome responsible for targeted protein degradation is much more abundant in *drmy1* than in WT (Supplemental Dataset 1). In concert, these results demonstrate that the machinery responsible for maintaining protein homeostasis is substantially dysregulated in *drmy1*.

A key signaling pathway that regulates protein homeostasis is TARGET OF RAPAMYCIN (TOR)^41,42^. TOR is a hub that integrates information from light, sugars, nutrient availability, etc., to promote growth-related processes, including ribosome biogenesis and translation, and to repress catabolic processes, including protein degradation by autophagy and the proteasome^33,34,43–45^. TOR directly regulates the translation of specific mRNAs by promoting the phosphorylation of proteins, including LARP1, eIF3h, RISP, eS6, and 4EBPs, that impact translation initiation, reinitiation, or elongation of mRNAs with specific features, such as 5’TOP motifs or short upstream open reading frames (uORFs) in the 5’ leaders of mRNAs^34,42,46–52^. TOR also indirectly increases overall protein synthesis rates by globally increasing ribosome levels. We therefore hypothesize that the overall decrease in ribosomal content and protein synthesis in *drmy1* may reflect altered TOR signaling. To test for signatures of transcriptomic changes that have been well defined in seedlings under TOR inhibition^34,53,54^, we performed RNA-seq on seedlings of WT, *drmy1*, WT treated with AZD-8055 (a potent TOR inhibitor), and mock-treated WT (Supplemental Dataset 3). We found that the *drmy1* mutation causes transcriptomic changes similar to TOR inhibition (Figure 1F). A significant portion of genes differentially expressed under TOR inhibition vs. mock were also differentially expressed in *drmy1* vs. WT (466/2044 = 22.8%; hypergeometric test, p = 4.7x10^-108^). Not only were these 466 genes differentially expressed in both situations, but also most of them were coherently downregulated or upregulated (439/466 = 94.2%, Chi-square test, p < 2.2x10^-16^; Figure 1G, S1I). Genes coherently downregulated in both situations were enriched in GO terms related to translation and ribosomes, and, most strikingly, a quarter of them were under the GO term “translation” (Figure 1F, S1J). These similar transcriptomic changes support our hypothesis that TOR activity is reduced in *drmy1*. To further test this hypothesis, we measured TOR activity in WT and *drmy1* by assaying the phosphorylation of its direct substrate, RIBOSOMAL PROTEIN eS6 KINASE (S6K)^55,56^. While the total protein level of S6K did not change in *drmy1*, we found that S6K phosphorylation drastically decreased, demonstrating reduced TOR activity (Figure 1H, I). Overall, these results are consistent with the idea that *drmy1* has reduced TOR activity—a main pathway controlling ribosomal abundance and translation—which causes reduced ribosomal content and global translation rate.

### Defects in TOR activity, ribosome, and translation disrupt robust sepal initiation

We next asked whether defects in TOR activity, ribosome, or translation have any effects on robust sepal initiation like the *drmy1* mutation does (Figure 2A, 2B; also see Zhu et al.^27^). In a WT bud, initiation is robust in that four sepal primordia of similar size form evenly spaced around the periphery of the floral meristem (Figure 2A, 2H). Angles between them vary little, i.e., they are all at around 90° angles from each other (Figure 2I, 2J). By contrast, in *drmy1* buds, three to five sepal primordia initiate and grow to different sizes (Figure 2B, 2H; also see Zhu et al.^27^). Sepal primordia in *drmy1* buds are generally unevenly spaced, and angles between them have a high coefficient of variation (CV) (Figure 2I, 2J). To determine whether defects in ribosomes can cause the same sepal initiation defects, we imaged three ribosomal mutants, *ul4z* (*rpl4a*), *ul4y*, and *ul18z* (*rpl5a*)^40^, each mutated in a gene encoding a ribosomal component that is also downregulated in *drmy1* at the transcript level (Figure S1C). The *ul4z* mutant bud shows reduced size of the inner sepal primordia relative to the outer sepal primordia (Figure 2C), and slightly more variable spatial distribution of sepal primordia (Figure 2J), although it always develops four sepal primordia (Figure 2H). This is a weaker phenotype than *drmy1* but has similar characteristics. The *ul4y* and *ul18z* mutants show great variability in the number and position of sepal primordia (Figure 2D, 2E, 2H, 2J), more similar to *drmy1*. We also crossed these ribosomal mutants with *drmy1* to study sepal variability in the double mutants (Figure S2A-H). In *drmy1 ul4z*, *drmy1 ul4y*, and *drmy1 ul18z/+*, on average, sepal initiation was as variable as in the *drmy1* single mutant (Figure S2I, S2J). However, there were buds with no outgrowth in the adaxial or lateral regions of the bud periphery (Figure S2B, S2E, S2G), buds with six sepal primordia (Figure S2C, S2F, S2H), and buds with two outer sepal primordia (Figure S2D, S2H), which were not seen in the *drmy1* single mutant. Note that we were unable to characterize the homozygous *drmy1 ul18z* double mutant because they were embryo-lethal (Figure S2K), further supporting the idea that ribosomal mutations enhance the phenotypic defects in *drmy1*.

**Figure 2.**
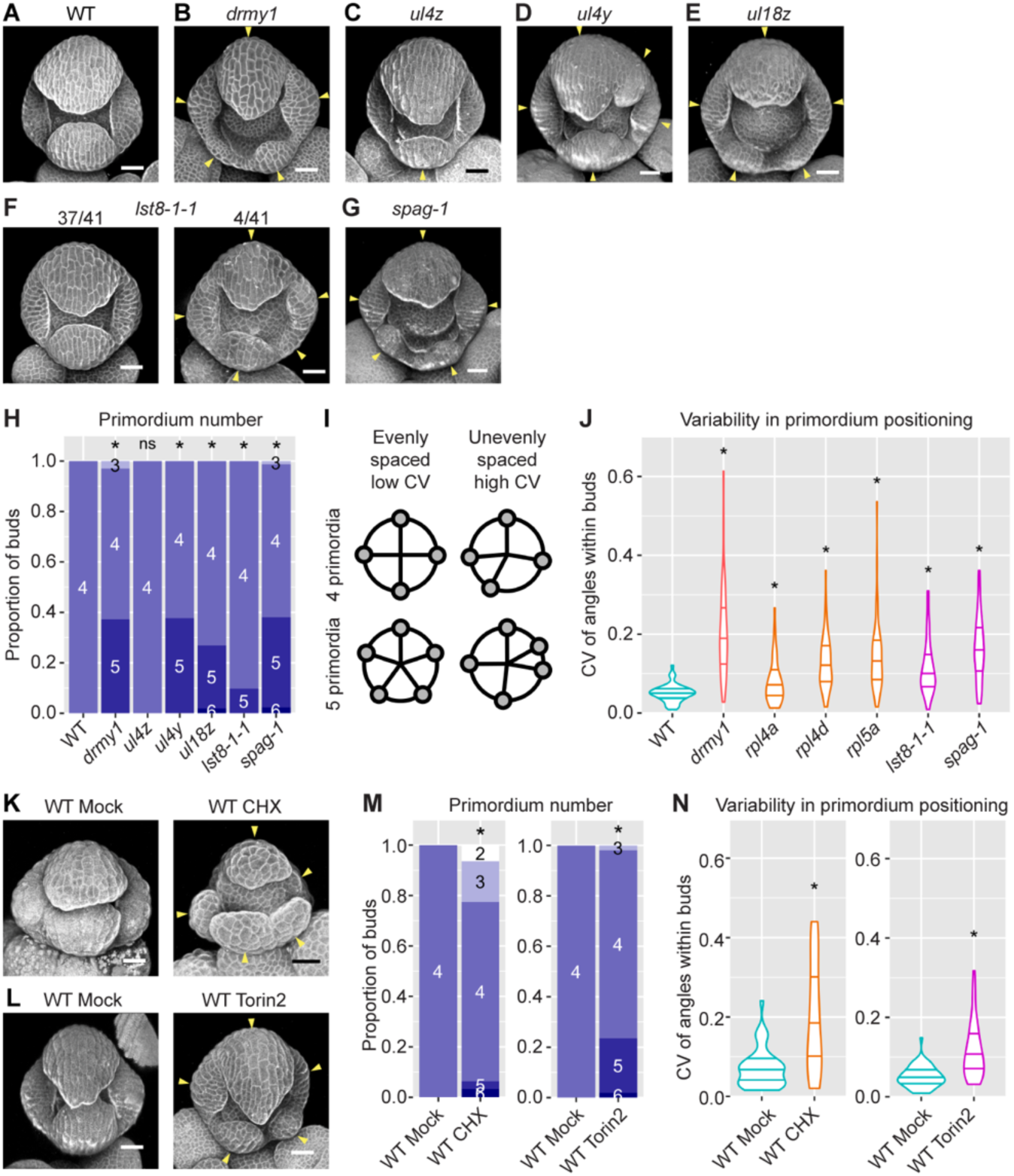
Defects in TOR activity, ribosome, and translation disrupt robust sepal initiation. **(A-G)** Representative images of stage 5 buds in WT (A), *drmy1* (B), *ul4z* (C), *ul4y* (D), *ul18z* (E), *lst8-1-1* (F), and *spaghetti-1* (G). Tissue morphology is visualized by either propidium iodide (a cell wall-staining dye) or a plasma membrane marker. Arrowheads indicate sepal primordia that are variable in number, position, and size. Note that *ul4z* flowers always develop four sepal primordia, although of different sizes; *lst8-1-1* occasionally (4/41, 9.8%) develops buds with more than four sepal primordia. **(H)** Quantification of sepal primordium number, comparing *drmy1* (n = 67 buds), *ul4z* (n = 52 buds), *ul4y* (n = 53 buds), *ul18z* (n = 52 buds), *lst8-1-1* (n = 41 buds), and *spaghetti-1* (n = 84 buds) with WT (n = 51 buds). Asterisks indicate statistically significant (p < 0.05) differences from WT in Fisher’s contingency table tests. **(I)** Illustration of robust versus variable positioning of sepal primordia. Primordia are considered robustly positioned if they are evenly distributed around the edge of the bud. Within each bud, angles between adjacent primordia with respect to the center of the bud are measured, and coefficient of variation (CV) is calculated. A bud with robustly positioned primordia would have similar angular values and a low CV value. A bud with variably positioned primordia would have very different angular values and a high CV value. **(J)** Quantification of variability in primordium positioning (CV) in the same buds as in (H), following illustration in (I). Asterisks indicate statistically significant (p < 0.05) differences from WT in Wilcoxon’s rank sum tests. **(K)** Representative images of buds from *in vitro*-cultured WT inflorescences treated with mock or 2 µM CHX for 9-10 days. Arrowheads indicate sepal primordia that are variable in number, position, and size. **(L)** Representative images of buds from WT plants treated with mock or 2 nmol Torin2 for 15 days. Arrowheads indicate sepal primordia that are variable in number, position, and size. **(M-N)** Quantification of sepal primordium number (M) and positional variability (N) similar to (H,J), comparing CHX-treated (n = 31 buds), CHX-mock (n = 42 buds), Torin2-treated (n = 51 buds) and Torin2-mock buds (n = 56 buds). Scale bars in all micrographs, 25 µm.

We then imaged mutants with reduced TOR activity to determine whether sepal initiation is also less robust. *lst8-1-1* is a T-DNA insertional mutant of the TOR complex component LST8-1^57^ and is weakly hypomorphic in TOR activity. We found that *lst8-1-1* shows variable sepal initiation in a small proportion of buds (4/41, 9.8%) (Figure 2F, 2H, 2J). The *spaghetti-1* mutant defective in TOR complex 1 (TORC1) assembly^58^ showed a level of variability comparable to the *drmy1* mutant and the ribosomal mutants *ul4y* and *ul18z* (Figure 2G, 2H, 2J). Mutants with more severe disruption of TOR activity are embryo lethal and could not be analyzed^58,59^. These results show that reduction in TOR activity can cause variability in sepal initiation, similar to *drmy1*.

To corroborate these findings, we directly inhibited translation by *in vitro* culture of dissected WT inflorescences on 2 µM cycloheximide (CHX, a chemical inhibitor of translation) for 9-10 days. This is a low concentration that does not completely block translation, as inflorescences were still alive after 10 days in this condition. Compared with mock, CHX-treated inflorescences develop buds that have 2 to 6 sepal primordia of variable sizes that are unevenly spaced around the bud periphery (Figure 2K, 2M, 2N). These phenotypes are stronger than *drmy1*. Similarly, we directly inhibited TOR activity by application of 2 nmol Torin2 to the growing shoot apex twice a day for 15 days, and we observed variable sepal initiation (Figure 2L, 2M, 2N). Overall, these data show that inhibition of TOR activity and translation can disrupt the robustness of sepal initiation, in terms of sepal primordium number, position, and size.

We previously showed that *drmy1* buds develop sepals of different sizes because of increased differences in the initiation timing of sepals within the same bud. The late-initiating primordia remain smaller throughout development. They end up as smaller sepals relative to those that initiated earlier, leaving gaps that expose the developing inner floral organs^27^. Moreover, different buds have different temporal patterns of sepal initiation, contributing to between-bud variability of sepal phenotype^27^. We asked whether TOR or ribosomal defects similarly disrupt the relative timing of sepal initiation, within-bud and between-bud. We live imaged WT and *ul4y* every six hours during sepal initiation and quantified the amount of time the bud takes to initiate the inner and lateral sepals after it initiates the outer sepal. In WT, after the initiation of the outer sepal, most buds initiate the inner sepal within 6 hours and the lateral sepals within 12 hours (Figure 3A, 3C; also see Zhu et al.^27^). In *ul4y*, the time differences between outer and inner sepal initiation and between outer and lateral sepal initiation are longer (Figure 3B, mean in Figure 3C). Moreover, these time differences are more variable across buds, indicating a loss of robustness in organ initiation timing (SD in Figure 3C). Similarly, we compared the sepal initiation timing in Torin2 vs mock-treated WT buds. While in most mock-treated buds, the inner and lateral sepals initiate within 12 hours after the outer sepal (Figure 3D, 3F), in Torin2-treated buds, sepals within the same bud initiates at vastly different times (Figure 3E, mean in Figure 3F), and the temporal pattern of sepal initiation is more variable across buds than mock (SD in Figure 3F). These results show that TOR and ribosomal defects can disrupt the precisely orchestrated initiation timing of sepal primordia.

**Figure 3.**
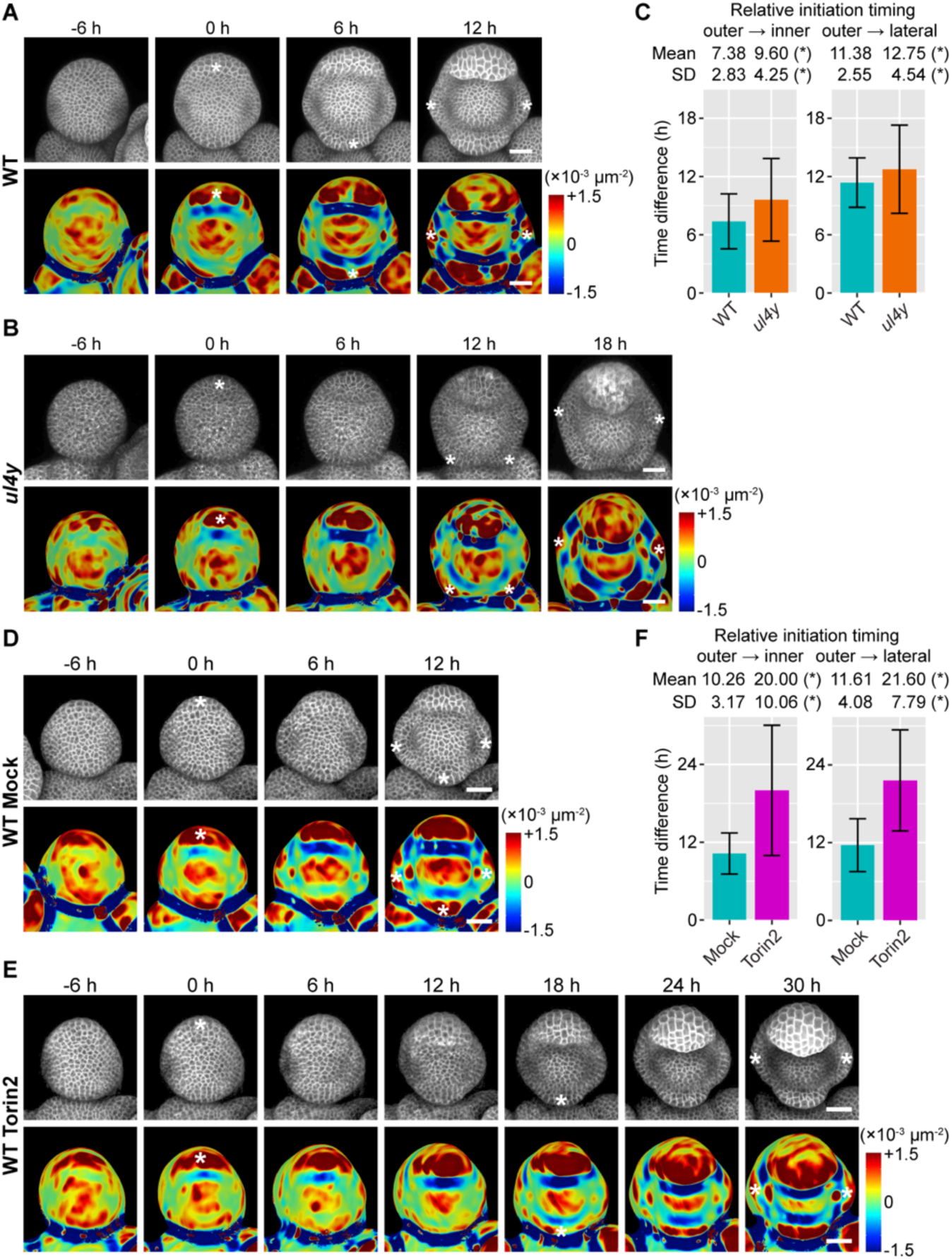
TOR and ribosomal defects cause variability in the timing of sepal initiation. **(A-C)** 6h-interval live imaging of the sepal initiation process in WT (A) and *ul4y* (B), which is quantified in (C). n = 48 buds for WT; n = 40 buds for *ul4y*. **(D-F)** 6h-interval live imaging of the sepal initiation process in buds from WT plants treated with mock or 2 nmol Torin2 twice a day for 15 days, which is quantified in (F). n = 31 buds for mock; n = 15 buds for Torin2. In **(A,B,D,E)**, top rows show the *35S::mCitrine-RCI2A* membrane marker, and bottom rows show Gaussian curvature heatmaps of the same image stacks. Asterisks indicate sepal initiation events, defined as a dark red band (primordium with positive curvature) separated from the floral meristem by a dark blue band (boundary with negative curvature) in the heatmap. Scale bars, 25 µm. In **(C,F)**, the amount of time between outer and inner sepal initiation (left) and between outer and lateral sepal initiation (right) were calculated for each bud. Bar plot shows mean ± SD which is also shown on top of each plot. Asterisks indicate statistically significant differences (p < 0.05) in Wilcoxon’s rank sum test (for mean) or Levene’s test (for SD).

Does the variability in initiation timing cause variable sizes and gaps in mature sepals, as in *drmy1* (Figure S3A, B, G, H; also see Zhu et al. ^27^)? We imaged the mature sepals of the ribosomal mutants *ul4z*, *ul4y*, *ul18z*, as well as the TOR component mutant *lst8-1-1*. Surprisingly, unlike *drmy1*, the sepals in *ul4z*, *ul4y*, *ul18z* enclose the inner floral organs perfectly, leaving no gaps, regardless of sepal number (Figure S3C-E). Small gaps still exist in buds of *lst8-1-1*, although sepal size differences appear greatly reduced (Figure S3F). Further dissection shows that in these mutants, sepals within the same flower are of similar sizes, although sepals from different flowers can be of vastly different sizes, most conspicuously for *lst8-1-1* (Figure S3I-N). This is unlike *drmy1*, where sepal size variability is equally high comparing sepals within the same flower or from different flowers (Figure S3H, S3M-N). Upon closer examination, while sepals initiating late in *drmy1* buds remain small, leaving a gap in the sepal whorl (Figure S3O-P), those in *ul4y* were able to catch up with the other sepals and close the gap (Figure S3Q). Our results suggest that there exists a size-coordinating mechanism independent of TOR or ribosomal functions that allows sepals within the same bud to reach the same mature length, and that this mechanism is disrupted in *drmy1*. Such a mechanism requires further investigation in the future.

### Inhibition of TOR activity and translation increase cytokinin signaling and disrupts the robust spatial pattern of auxin and cytokinin signaling

Auxin and cytokinin are two important plant hormones critical to many aspects of plant development^30–32^, and there is accumulating evidence that they act synergistically in the shoot apical meristem to promote lateral organ initiation^16,60,61^. We previously showed that, in a WT floral meristem prior to sepal initiation, auxin and cytokinin signaling are concentrated at the four incipient primordia, which is required for robust sepal initiation from these regions (Figure 4A, S4A; Zhu et al.^27^). In the *drmy1* mutant, cytokinin signaling becomes stronger and diffuse around the bud periphery (Figure 4A-B). Auxin signaling also becomes more diffuse, forming irregular auxin maxima that are less focused than those in WT, except at the incipient outer sepal where it remains robust (Figure 4A, S4B; Zhu et al.^27^). These changes in hormone signaling correlate with variable sepal initiation (Figure S4B)^27^. We wondered whether ribosomal mutations have similar effects on auxin and cytokinin signaling. To this end, we imaged the auxin signaling reporter *DR5::3xVENUS-N7* and the cytokinin signaling reporter *TCS::GFP* in floral meristems of the ribosomal mutant *ul4y*. Both reporters lose their robust spatial pattern except in the incipient outer sepal (Figure 4A, S4C). The hormone signaling patterns were quantified by circular histogram analysis (see Methods for details). For each of DR5 and TCS, WT buds showed four clear peaks ∼90 degrees apart from each other, with very little signal in between, whereas in *drmy1* and *ul4y*, peaks were barely seen except at the incipient outer sepal (at 45 degrees), and there was greater noise and variation all around the bud (Figure 4C-D). Diffuse bands of auxin signaling that typically occurs in the adaxial or lateral periphery of *drmy1* and *ul4y* buds (Figure S4B and S4C, brackets) can later resolve into several distinct auxin maxima of various intensity and at various positions, correlated with the initiation of sepal primordia of various sizes at these same positions (Figure S4B and S4C, red arrowheads).

**Figure 4.**
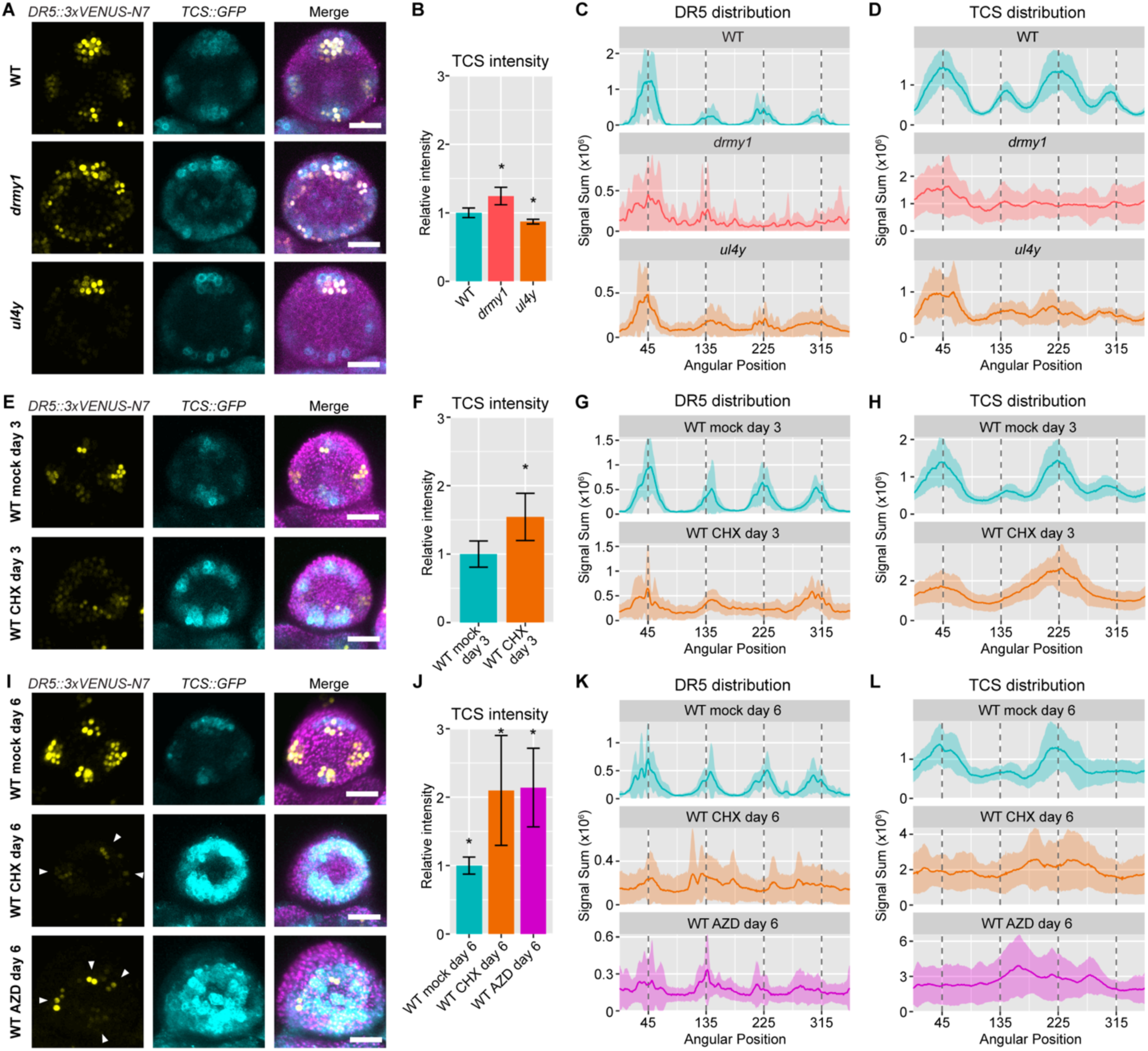
Inhibition of TOR activity and translation cause variability in auxin and cytokinin signaling. **(A-D)** The ribosomal mutant *ul4y* loses robustness in auxin and cytokinin signaling. (A) Representative images of late stage 2 buds of WT, *drmy1*, and *ul4y*, showing the auxin signaling reporter *DR5::3xVENUS-N7* in yellow, the cytokinin signaling reporter *TCS::GFP* in cyan, and both merged with Chlorophyll (in WT) or *UBQ10::mCherry-RCI2A* (in *drmy1* and *ul4y*) in magenta. (B) Quantification of TCS intensity (integrated density divided by area) from maximum intensity projection images, normalized to mean of WT. Shown are mean ± SD. Asterisks show statistically significant differences from WT in two-tailed Student’s t-tests (*drmy1*, p = 2.1×10^-6^; *ul4y*, p = 3.4×10^-5^). (C) Circular histogram of DR5 signal distribution. Each bud was divided into 360 sectors of 1° each. Within each sector, DR5 signal measured in pixel intensity units (0-255 range) was summed. This sum was plotted along the x-axis starting from the sector at 1:30 position (between the incipient outer sepal and incipient right sepal) going counterclockwise. I.e., in WT, the outer sepal is near 45°, the inner sepal near 225°, and the lateral sepals near 45° and 135° (vertical dotted lines). The mean was plotted as a solid line, and mean ± SD was plotted as a shaded area.(D) Circular histogram of TCS signal distribution. Sample size for (A-D): WT, n = 12 buds; *drmy1*, n = 15 buds; *ul4y*, n = 10 buds. **(E-H)** 3 days of translation inhibition causes increased and diffuse cytokinin signaling, and diffuse auxin signaling. (E) Representative images of late stage 2 WT buds treated *in vitro* with mock or 2 µM CHX for 3 days. Shown are *DR5::3xVENUS-N7* in yellow, *TCS::GFP* in cyan, and both merged with Chlorophyll in magenta. (F) Quantification of TCS intensity from maximum intensity projection images, normalized to mean of WT mock day 3. Shown are mean ± SD. Asterisk shows statistically significant difference in a two-tailed Student’s t-test (p = 2.0×10^-4^). (G) Circular histogram of DR5 signal distribution. (H) Circular histogram of TCS signal distribution. Sample size for (E-H): WT mock day 3, n = 10 buds; WT CHX day 3, n = 12 buds. **(I-L)** 6 days of TOR or translation inhibition causes increased and diffuse cytokinin signaling, and randomly positioned auxin signaling maxima. (I) Representative images of late stage 2 WT buds treated *in vitro* with mock, 2 µM CHX, or 2 µM AZD for 6 days. Shown are *DR5::3xVENUS-N7* in yellow, *TCS::GFP* in cyan, and both merged with Chlorophyll in magenta. Arrowheads point to randomly positioned auxin maxima. (J) Quantification of TCS intensity from maximum intensity projection images, normalized to mean of WT mock day 6. Shown are mean ± SD. Asterisks show statistically significant differences from mock in two-tailed Student’s t-tests (CHX, p = 1.0×10^-3^; AZD, p = 1.2×10^-4^). (K) Circular histogram of DR5 signal distribution. (L) Circular histogram of TCS signal distribution. Sample size for (I-L): WT mock day 6, n = 12 buds; WT CHX day 6, n = 11 buds; WT AZD day 6, n = 10 buds. Scale bars in all micrographs, 25 µm.

We also tested whether drug treatments that inhibit TOR activity or translation can disrupt the robust hormone patterning. Buds treated *in vitro* with the translation inhibitor CHX (2 µM) for 3 days showed a 50% increase in cytokinin signaling, and both auxin and cytokinin signaling became diffuse around the bud periphery (Figure 4E-H). By day 6, cytokinin signaling was still diffuse all around, and increased to more than two-fold relative to mock (Figure 4I, 4J, 4L). Auxin signaling formed maxima of variable number at variable positions (Figure 4I arrowheads, 4K), correlated with the variable initiation of sepal outgrowth at these positions (Figure S4D-E). Similar changes occurred in buds treated *in vitro* with the TOR inhibitor AZD-8055 (2 µM) for 6 days (Figure 4I-L). For both CHX and AZD-8055, the disruptions of hormone signaling are similar to *drmy1*. *In vivo* treatment using another TOR inhibitor Torin2 for 15 days increased cytokinin signaling by 70%, although it did not make auxin and cytokinin signaling more diffuse (Figure S4F-I). Overall, these results show that defects in TOR activity and translation increase cytokinin signaling, and disrupt the precise spatial patterning of cytokinin and auxin signaling required for robust sepal initiation.

### An increase in cytokinin signaling is necessary and sufficient for variable auxin signaling and sepal initiation under translation inhibition

Auxin is a critical hormone in organogenesis^62,63^. As shown above, variable patterning of auxin signaling correlates with variable sepal initiation during inhibition of TOR activity and translation. We wondered what caused auxin to lose its robust patterning under such conditions. It was previously reported that the ribosomal mutants *ul4y*, *ul18z*, and *el24y* have reduced protein levels of AUXIN RESPONSE FACTOR (ARF) 3, 5, and 7^64–66^, key transcription factors that mediate the auxin signaling response^67^. The transcripts of these ARFs contain upstream open reading frames (uORFs), requiring translation reinitiation to translate their main open reading frames^68,69^, a process defective in the ribosomal mutants *ul4y*, *ul18z*, and *el24y*^64–66^. We therefore hypothesized that *drmy1* loses robust auxin signaling pattern because of reduced translation of uORF-containing transcripts, including those of certain ARFs. To begin, we utilized our transcriptomics and proteomics data, and considered that the protein-transcript ratio of a gene should reflect its level of translation, among other factors such as protein stability. Therefore, following our hypothesis, genes containing uORFs should, in general, have a lower protein-transcript ratio in *drmy1* than in WT. We calculated the difference of this ratio between *drmy1* and WT for all 5,086 gene-protein pairs in our inflorescence dataset, and compared the ratio against the number of uORFs in each transcript (Figure S5A; uORF data from von Arnim et al.^69^). We found a small but significant decrease in the protein-transcript ratio in *drmy1* for the 724 genes containing at least 2 uORFs in their transcripts, supporting the hypothesis that *drmy1* has reduced translation reinitiation for uORF-containing transcripts, just like *ul4y*, *ul18z*, and *el24y*^64–66^. Then, we examined whether the translation reinitiation of uORF-containing ARFs are indeed reduced in the *drmy1* mutant. We selected *ARF3/ETTIN*, *ARF5/MONOPTEROS*, and *ARF6*, which have 2, 6, and 6 uORFs respectively, and as controls, *ARF8* and *ARF10* which do not contain uORFs. None of these *ARF*s were differentially expressed in *drmy1* at the transcript level, except *ARF10* which was slightly upregulated (Figure S5B). We utilized promoter-fluorescent protein fusion reporters (Figure S5C) which have the same uORFs in the promoter region as the corresponding ARF genes if the genes have them. These reporters reflect transcriptional and uORF-mediated translational regulation. *pARF3::N3xGFP*, *pARF5::ER-EYFP-HDEL*, and *pARF6::N3xGFP* contain uORFs and thus, following our hypothesis, are expected to drastically decrease in fluorescence intensity in *drmy1* compared to WT. *pARF8::N3xGFP* and *pARF10::N3xGFP* do not have uORFs and are thus expected to have comparable or higher fluorescence intensity in *drmy1*. Surprisingly, we saw no correlation between the presence of uORFs and decrease in fluorescent intensity in *drmy1* (Figure S5C-D). While it might arise from additional layers of regulation on these ARFs, this result suggests that the decrease in translation reinitiation of uORF-containing ARFs is not the main factor explaining the loss of robust auxin signaling pattern in *drmy1*.

It was previously reported that external application of cytokinin increases auxin biosynthesis in actively growing tissue including the shoot apex, young leaves, and roots^70^, and cytokinin application also changes the expression and polarity of PIN-FORMED (PIN) polar auxin transport carriers^71,72^. We previously noticed that external application of 6-benzylaminopurine (BAP), a synthetic cytokinin, induced additional convergence points of PIN1 and increased variability in auxin signaling, causing variability in sepal initiation (Zhu et al.^27^, in this reference see Fig. 4e, Extended Data Fig. 7e and 7f). Here, we confirmed this observation by circular histogram analysis (Figure 5A-D). While the mock-treated WT buds showed four clear peaks of DR5 signal with very little signal in between (Figure 5A-B), those treated with 5 µM BAP showed a less robust spatial pattern, with less distinguishable peaks and larger variation all around the bud (Figure 5C-D). Thus, excessive cytokinin is sufficient for the variable spatial pattern of auxin signaling.

**Figure 5.**
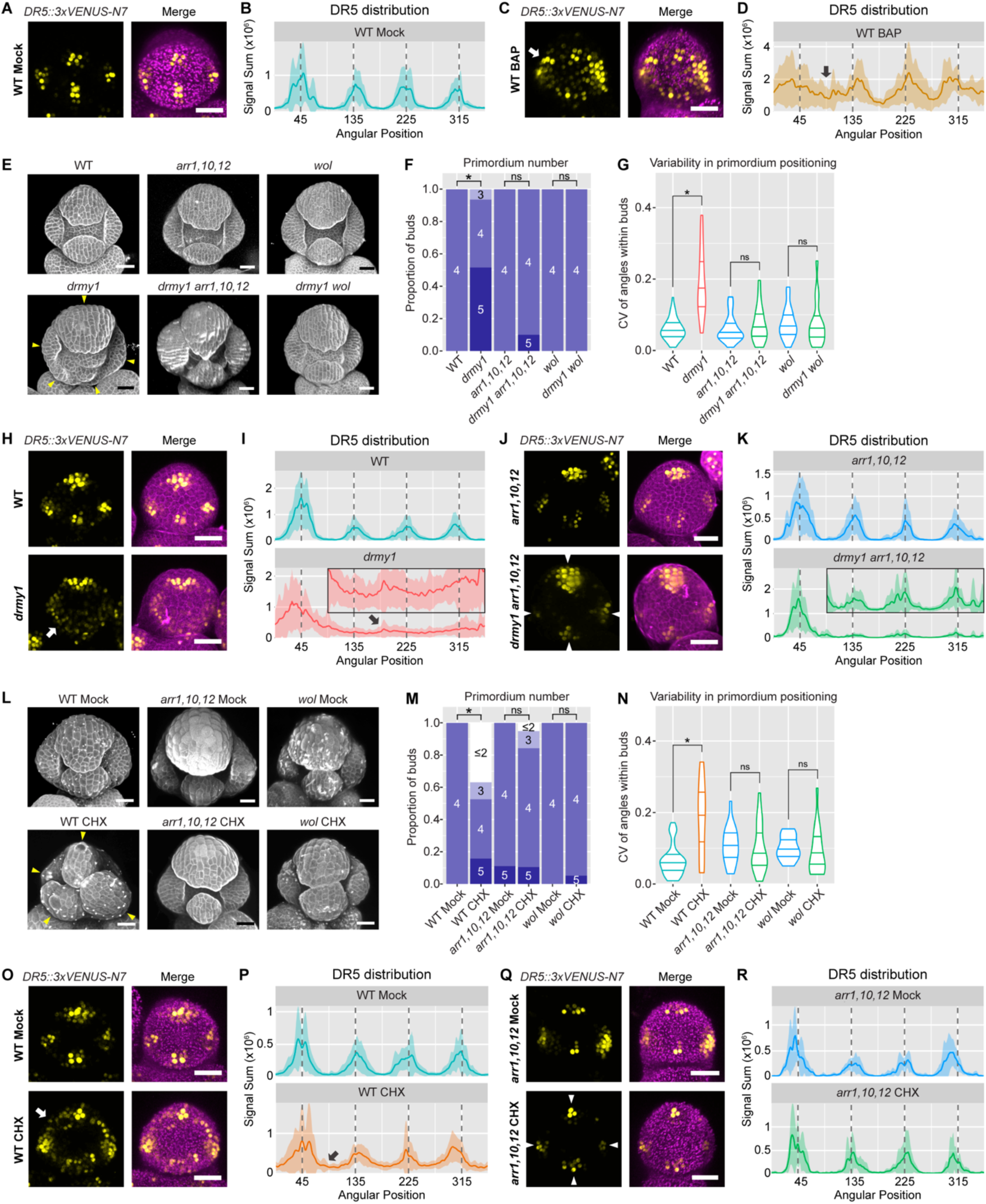
Cytokinin signaling is required for increased variability in auxin signaling and sepal initiation under translation inhibition. **(A-D)** Cytokinin treatment makes auxin signaling diffuse. Shown are late stage 2 WT buds under mock (A,B) or 5 µM cytokinin (BAP) treatment (C,D) for 4 days. (A,C) Auxin signaling reporter *DR5* in yellow, and *DR5* merged with Chlorophyll in magenta. (B,D) Circular histograms of the *DR5* signal, showing mean (solid line) and mean ± SD (shaded area). Arrows point to DR5 signal in variable positions. Sample size: WT Mock n = 10, WT BAP n = 10. Also see Zhu et al. (2020), in this reference see Extended Data Figure 7e. **(E-G)** Cytokinin signaling is required for variable sepal initiation in *drmy1*. (E) Stage 5 buds. Sepal primordia in *drmy1* are variable (arrowheads), which does not occur in *drmy1 arr1,10,12* and *drmy1 wol* mutants. (F,G) Quantification of sepal primordium number (F) and positional variability (G), comparing WT (n = 58) with *drmy1* (n = 31), *arr1,10,12* (n = 24) with *drmy1 arr1,10,12* (n = 20), and *wol* (n = 36) with *drmy1 wol* (n = 39). Asterisks indicate statistically significant (p < 0.05) differences in Fisher’s contingency table tests (F) and Wilcoxon’s rank sum tests (G) respectively. **(H-K)** Cytokinin signaling is required for variable patterning of auxin signaling in *drmy1*. Shown are late stage 2 buds of WT vs *drmy1* (H,I), and *arr1,10,12* vs *drmy1 arr1,10,12* (J,K). (H,J) Auxin signaling reporter *DR5* in yellow, and *DR5* merged with propidium iodide in magenta. Arrows point to diffuse DR5 signal in variable positions of the *drmy1* bud. Arrowheads show four robust DR5 maxima in the *drmy1 arr1,10,12* bud. (I,K) Circular histograms of the *DR5* signal, showing mean (solid line) and mean ± SD (shaded area). For ease of visualization, circular histograms of *drmy1* and *drmy1 arr1,10,12* between 90 and 360 degrees are enlarged and shown as insets (y-axis range 0-0.4). Note the presence of DR5 signal in inter-sepal regions in *drmy1* (black arrow) which is largely suppressed in *drmy1 arr1,10,12*. Sample size: WT n = 19, *drmy1* n = 16, *arr1,10,12* n = 13, *drmy1 arr1,10,12* n = 9. **(L-N)** Cytokinin signaling is required for variable sepal initiation under translation inhibition. (L) Stage 6 buds of WT, *arr1,10,12*, and *wol*, treated with Mock or 2 µM CHX for 10 days. WT initiates sepal primordia at variable positions when treated with CHX (arrowheads), which does not occur in *arr1,10,12* and *wol*. (M,N) Quantification of sepal primordium number (M) and positional variability (N), comparing mock and CHX within each genotype. Sample size: WT Mock n = 29, WT CHX n = 19, *arr1,10,12* Mock n = 18, *arr1,10,12* CHX n = 19, *wol* Mock n = 15, *wol* CHX n = 19. Asterisks indicate statistically significant (p < 0.05) differences in Fisher’s contingency table tests (M) and Wilcoxon’s rank sum tests (N) respectively. **(O-R)** Cytokinin signaling is required for diffuse auxin signaling under translation inhibition. Shown are late stage 2 buds of WT (O,P) and *arr1,10,12* (Q,R), treated with Mock or 2 µM CHX for 3 days. (O,Q) Auxin signaling reporter *DR5* in yellow, and *DR5* merged with Chlorophyll in magenta. Arrows point to diffuse DR5 signal in variable positions in CHX-treated WT. Arrowheads show four robust DR5 maxima in CHX-treated *arr1,10,12*. (P,R) Circular histograms of the *DR5* signal, showing mean (solid line) and mean ± SD (shaded area). Sample size: WT Mock n = 17, WT CHX n = 18, *arr1,10,12* Mock n = 7, *arr1,10,12* CHX n = 7. Scale bars in all micrographs, 25 µm.

We then wondered whether an increase in cytokinin signaling (Figure 4) is the cause of variable pattern of auxin signaling under translation-limited conditions such as *drmy1*. To test this hypothesis, we crossed *drmy1* containing the DR5 reporter with a triple mutant of *ARABIDOPSIS RESPONSE REGULATOR* (*ARR*) *1*, *10*, and *12*, the three most highly expressed B-type ARRs in our RNA-seq (Supplementary Dataset 1) which are crucial for the activation of cytokinin-responsive genes^73^. While buds of *arr1,10,12* did not show apparent phenotypic differences from WT, the quadruple mutant *drmy1 arr1,10,12* largely rescued the *drmy1* phenotype, with much less variability in sepal number and position (Figure 5E-G). While mature buds of *drmy1* have sepals of variable sizes, leaving gaps and exposing the inner floral organs (Figure S6D vs. S6A-C), those of *drmy1 arr1,10,12* have sepals of robust sizes that are able to close (Figure S6E). Likewise, mutation in a cytokinin receptor *WOODEN LEG* (*WOL*)/*ARABIDOPSIS HISTIDINE KINASE 4* (*AHK4*) showed a similar rescue of the *drmy1* sepal phenotype (Figure 5E-G, S6F). While the auxin signaling reporter DR5 was diffuse and variable in *drmy1* except in the incipient outer sepal (Figure 5H-I), in *drmy1 arr1,10,12*, it was focused in all the four incipient sepals that were robustly positioned, although the signal intensity in the incipient outer sepal was much higher than others (Figure 5J-K). These results indicate that cytokinin signaling is required for the increased variability in auxin signaling pattern and sepal initiation in *drmy1*.

Furthermore, we wanted to test whether cytokinin signaling is required for variability in more general conditions where translation is inhibited. The translation inhibitor CHX disrupted robustness in auxin signaling and sepal initiation in WT (Figure 2K, 4E, 4I), and we tested whether these effects are still present in *arr1,10,12* and *wol* mutants. We found that, unlike WT, sepal initiation remained mostly robust in *arr1,10,12* and *wol* after ten days of 2 µM CHX treatment (Figure 5L-N). While DR5 in WT became diffuse and occurred in variable positions after three days of CHX treatment (Figure 5O-P, arrow), DR5 in *arr1,10,12* remained robust and concentrated at the four incipient sepal primordia (Figure 5Q-R). These results suggest that elevated cytokinin signaling level is the primary cause for variability in auxin patterning under translation-inhibited conditions. Thus, in WT, maintaining a low level and focused cytokinin signaling is crucial for robust auxin patterning and sepal initiation.

### Upregulation of cytokinin signaling is required to sustain translation and fitness in *drmy1*

Under translation-inhibited conditions, why does the plant upregulate cytokinin signaling at the cost of robust morphogenesis? Previous studies revealed that cytokinin signaling can stimulate translation^74–78^, by increasing transcription or protein abundance of ribosomal components or biogenesis factors^79–81^ and modification of initiation and elongation factors^82^. We therefore hypothesized that an increase in cytokinin signaling under translation-inhibited conditions (such as *drmy1*) sustains a survivable rate of translation in a feedback loop. We first validated that, under our growth conditions, an increase in cytokinin signaling (*arr1 35S::ARR1*) is sufficient to increase global translation (Figure 6A; also see Karunadasa et al.^74^) in 14-day-old seedlings. We then tested whether cytokinin signaling is required to sustain global translation (Figure 6B-C). Compared to WT, the cytokinin receptor single mutant *wol* has a mild reduction in global translation rate at day 8 and a ∼50% reduction at day 14. The *drmy1* single mutant shows drastically reduced global translation rate at day 8, but by day 14, global translation rate in *drmy1* increased and matched WT. In the *drmy1 wol* double mutant, however, translation rate was unable to recover at day 14 and remained lower than *drmy1*. Our data suggest that, in *drmy1* plants which has reduced TOR activity and ribosomal level (Figure 1), the upregulated cytokinin signaling is required to sustain global translation at nearly WT levels.

**Figure 6.**
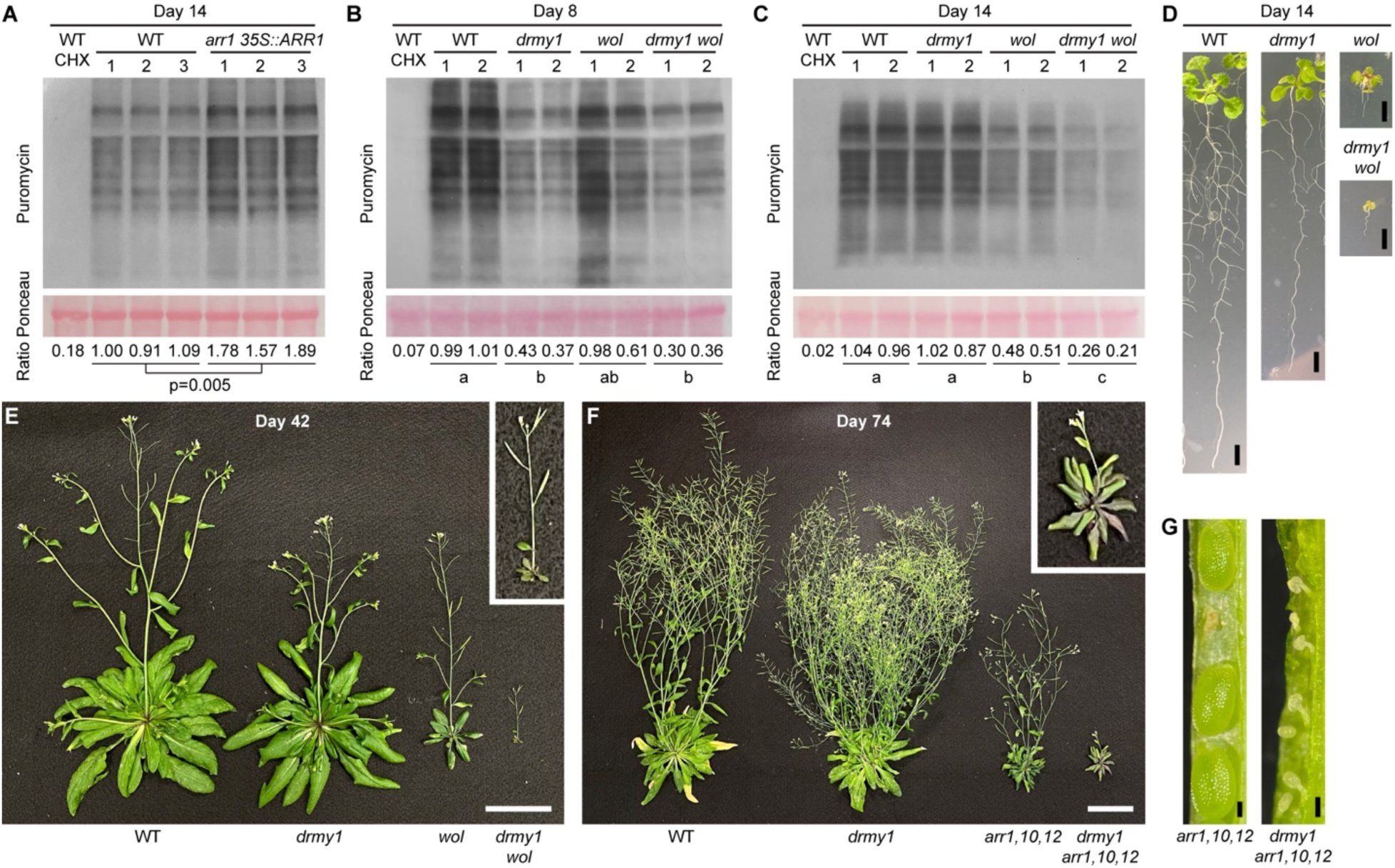
Upregulation of cytokinin signaling is required to maintain translation and fitness in *drmy1*. **(A)** Puromycin labeling of WT seedlings with 4 h CHX pre-treatment (control), and three biological replicates each of WT and *arr1 35S::ARR1* seedlings with 4 h mock pre-treatment. All seedlings are 14 days old. RuBisCO large subunit in Ponceau S-stained membrane is shown as a loading control. Signal ratio between puromycin and Ponceau S, normalized to mean of WT, is show on the bottom. P-value is from a two-sided Student’s t-test. Also see Karunadasa et al. (2020). **(B,C)** Puromycin labeling of WT seedlings with 4 h CHX pre-treatment (control), and two biological replicates of WT, *drmy1*, *wol*, and *drmy1 wol* seedlings with 4 h mock pre-treatment. Seedlings are 8 days old in (B) and 14 days old in (C). RuBisCO large subunit in Ponceau S-stained membrane is shown as a loading control. Letters show compact letter display of a Tukey’s all-pair comparison in a one-way ANOVA model. **(D)** Representative 14 days old seedling images of WT, *drmy1*, *wol*, and *drmy1 wol* used in (C). Notice that *drmy1 wol* is very small and pale. Scale bars, 5 mm. **(E)** Representative aerial part images of 42 days old plants of WT, *drmy1*, *wol*, and *drmy1 wol*. Inset shows the zoomed-in *drmy1 wol* plant, which has a tiny rosette and a short inflorescence. Scale bars, 5 cm. See also Figure S6F. **(F)** Representative aerial part images of 74 days old plants of WT, *drmy1*, *arr1*,*10*,*12*, and *drmy1 arr1*,*10*,*12*. Inset shows the zoomed-in *drmy1 arr1*,*10*,*12* plant, which has pale leaves accumulating anthocyanin and a short inflorescence. Scale bars, 5 cm. See also Figure S6E. **(G)** Dissected siliques of *arr1,10,12* (left) and *drmy1 arr1,10,12* (right) showing developing seeds. Notice that while *arr1,10,12* occasionally have aborted seeds, all seeds in the *drmy1 arr1,10,12* silique were aborted. Scale bars, 0.2 mm.

We then hypothesized that an upregulation of cytokinin signaling in plants with reduced TOR activity and ribosomal content such as *drmy1* and the consequent restoration of global translation would provide fitness benefits. Thus, we expect that removal of the cytokinin receptor WOL from *drmy1* and the consequent failure to sustain global translation should affect plant vitality and reproduction. We found that at day 14, the *drmy1* single mutant is slightly smaller than WT. In contrast, in the absence of WOL, growth of *drmy1 wol* plants were extremely retarded compared to *wol*, with tiny and chlorotic cotyledons and true leaves (Figure 6D). In older plants, the *drmy1* single mutant has similar rosette size and slightly shorter inflorescences compared to WT. In contrast, in the absence of WOL, *drmy1 wol* produced tiny rosettes and stunted inflorescences with a few chlorotic buds that develop into small, short siliques (Figure 6E, S6F). Similarly, when B-type ARRs are mutated, the growth of the *drmy1 arr1,10,12* quadruple mutant is much retarded compared to *arr1,10,12*. They produced slightly chlorotic and anthocyanin-rich rosettes, a tiny inflorescence composed of very few buds (Figure 6F, S6E) and, in the end, siliques in which all seeds had aborted (Figure 6G). Overall, these results show that *drmy1* requires the cytokinin signaling pathway for normal growth and reproduction. While it remains possible that unknown effects of the cytokinin signaling pathway other than promoting translation are critical for the proper growth of *drmy1*, our results are consistent with our hypothesis that the upregulation of global translation (Figure 6A) by increased cytokinin signaling (Figure 4) maintains a survivable level of protein synthesis in plants with reduced ribosomal content such as *drmy1* (Figure 1C).

### TOR and translation inhibition decreases the protein level of cytokinin signaling inhibitors ARR7 and AHP6

What causes cytokinin signaling to increase in plants with reduced TOR activity and translation (Figure 4)? It was previously shown that *cis*-type cytokinins can be synthesized from tRNAs by the tRNA isopentenyltransferases (IPTs), IPT2 and IPT9^83^. We hypothesized that the decrease in global translation may increase the availability of tRNAs as substrates for cytokinin biosynthesis, increasing the level of cytokinins. To test this idea, we extracted cytokinins from induced inflorescences of WT and *drmy1* in *ap1 cal AP1-GR* background (Figure S1A). We measured the level of three cytokinin bases, *trans*-Zeatin (tZ), *cis*-Zeatin (cZ), and isopentenyladenine (iP), and their corresponding nucleosides (tZR, cZR, and iPR), using liquid chromatography-mass spectrometry. Surprisingly, we found no significant difference in their levels between WT and *drmy1*, and notably, the amount of *cis*-Zeatin was barely detectable in all samples (Figure S7A). This suggests that the increase in *cis*-type cytokinin synthesis is not the mechanism underlying the increase in cytokinin signaling under our translation-inhibited conditions.

We then considered the effects that a decrease in translation rate might have on the protein components of the cytokinin signaling pathway. In particular, A-type *ARR*s, which encode inhibitors of cytokinin signaling^84–86^, are rapidly induced upon cytokinin application and serve to dampen cytokinin response in the tissue^87–89^. Likewise, *AHP6* is highly expressed in lateral organ primordia downstream of auxin signaling, which non-cell autonomously represses and restricts cytokinin signaling to robust spatial patterns^16^. The rapid synthesis of the A-type ARR and AHP6 proteins may be crucial for maintaining the homeostasis of cytokinin signaling during developmental processes. We therefore hypothesized that, during hormone patterning prior to sepal initiation, translation defects in *drmy1* cause reduced synthesis of these proteins, decreasing them to a level insufficient to repress cytokinin signaling (Figure 7A).

**Figure 7.**
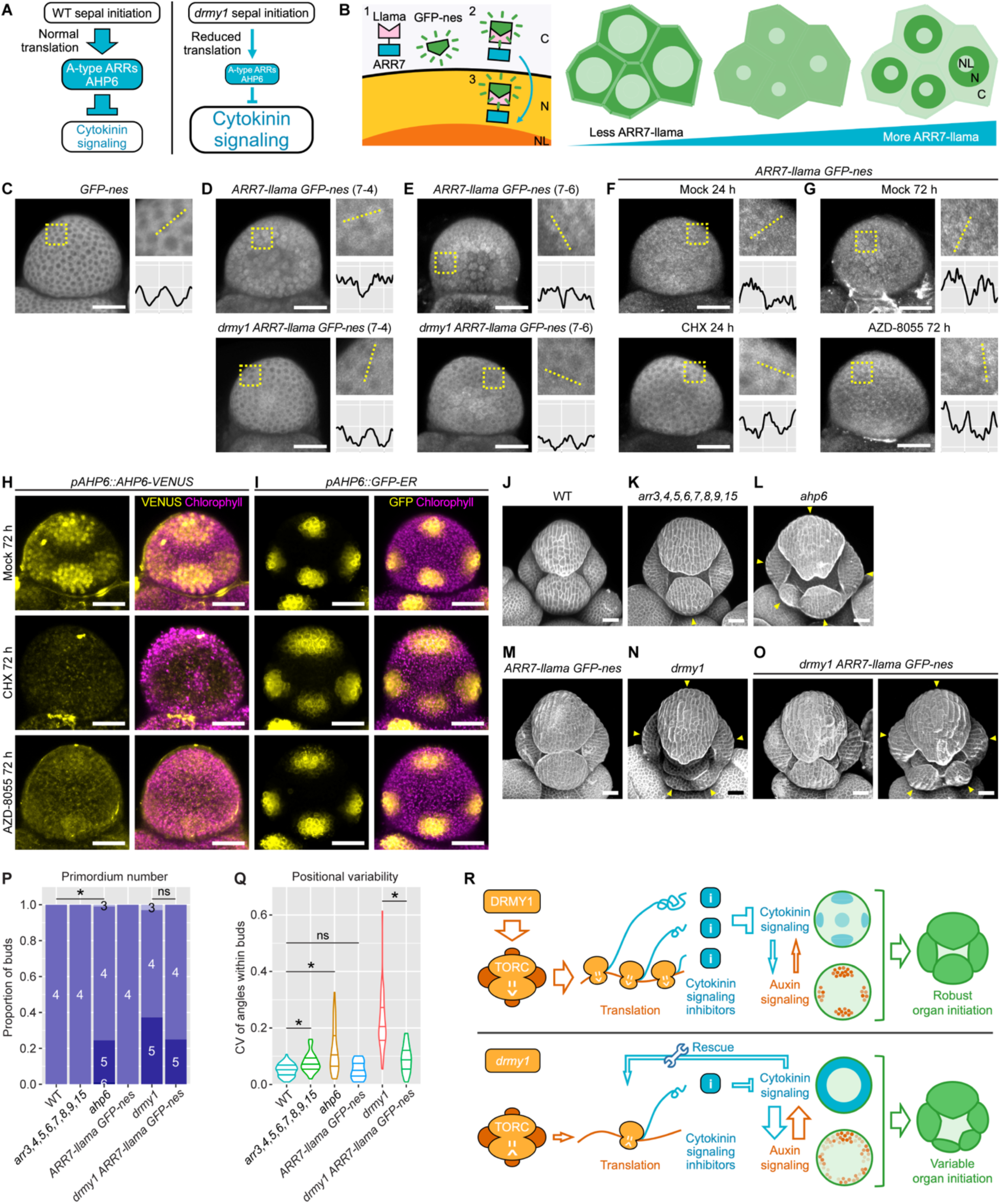
ARR7 and AHP6 protein levels are reduced upon inhibition of TOR and translation. **(A)** The hypothesis. During hormone patterning prior to sepal initiation in the WT floral meristem, A-type ARR and AHP6 proteins are rapidly produced to dampen cytokinin signaling to a normal level. In *drmy1*, reduced protein synthesis causes reduced levels of these cytokinin signaling inhibitor proteins, resulting in an upregulation of cytokinin signaling. **(B)** Illustration of Llama Tag. Plants were co-transformed with *ARR7-llama* (*pARR7::ARR7-linker-llama-ARR7ter*) and *GFP-nes* (*pUBQ10::sfGFP-nes-UBQ3ter*). Without ARR7-llama, GFP localizes to the cytosol due to the nuclear export sequence (nes). ARR7-llama is produced in the cytoplasm, C, and translocates into the nucleus, N. The Llama Tag on ARR7-llama binds to GFP and drags GFP into the nucleus (note that from our observation it is excluded from the nucleolus, NL). Thus, at low ARR7-llama levels, GFP signal is mainly in the cytoplasm. At intermediate ARR7-llama levels, GFP is at comparable levels between the cytoplasm and the nucleus, and no clear pattern can be seen. At high ARR7-llama levels, GFP is mainly seen in the nucleus. **(C)** A *GFP-nes* bud showing localization of the GFP signal to the cytoplasm. **(D,E)** GFP images of buds from two independent transgenic lines of *ARR7-llama GFP-nes*, 7-4 (D) and 7-6 (E), of WT (top) vs *drmy1* (bottom) genotypes. Images are representative of n = 17 (line 7-4, WT), n = 40 (line 7-4, *drmy1*), n = 9 (line 7-6, WT), and n = 6 (line 7-6, *drmy1*) buds. Note that GFP is more cytoplasm-localized in *drmy1* than WT, indicating reduced ARR7-llama protein level. **(F)** GFP images of WT *ARR7-llama GFP-nes* buds treated with mock (top) or 2 µM CHX (bottom) for 24 hours. The mock image is representative of n = 20 buds from three independent lines. The CHX image is representative of n = 19 buds from these same lines. **(G)** GFP images of WT *ARR7-llama GFP-nes* buds treated with mock (top) or 2 µM AZD-8055 (bottom) for 72 hours. The mock image is representative of n = 13 buds from two independent lines. The AZD-8055 image is representative of n = 11 buds from these same lines. For (C-G), each image was brightened to reveal subcellular localization patterns of GFP. A square region taken from the image containing 5-10 cells is enlarged and shown on the top right. Within the square, GFP intensity was quantified along the dotted line and plotted on the bottom right. X-axis, pixels (range 0-238). Y-axis, GFP intensity (smoothened by taking the average intensity of 11-pixel neighborhoods; range 90-210 in gray value). **(H-I)** Response of the AHP6 protein reporter (H) and transcriptional reporter (I) to mock, CHX, and AZD-8055 treatments for 72 hours. For (H), images are representative of n = 29 (mock), n = 29 (CHX), and n = 34 (AZD-8055) buds in three experiments. For (I), images are representative of n = 11 (mock), n = 9 (CHX), and n = 12 (AZD-8055) buds in two experiments. **(J-Q)** Reduction of A-type ARR and AHP6 protein levels contribute to the variability in sepal initiation. (J-O) Stage 5-6 buds of indicated genotype stained with propidium iodide. Arrowheads indicate sepal primordia that are variable in number, position, and/or size. Note that the *arr3,4,5,6,7,8,9,15* bud has an inner sepal that is slightly smaller than its outer sepal and positioned slightly right-skewed (K). The *ahp6* bud develops five sepal primordia of variable sizes and unevenly positioned (L). The *ARR7-llama GFP-nes* constructs partially rescue the *drmy1* phenotype in some buds (O, left) but not others (O, right). (P) Quantification of sepal primordium number. Asterisk indicates statistically significance difference in a Fisher’s exact test (WT vs *ahp6*, p = 3.026×10^-7^; *drmy1* vs *drmy1 ARR7-llama GFP-nes*, p = 0.4389). (Q) Quantification of variability in sepal primordium position. Asterisk indicates statistically significant difference in a Wilcoxon rank sum test (WT vs. arr3,4,5,6,7,8,9,15, p = 2.948×10^-4^; WT vs *ahp6*, p = 2.137×10^-11^; WT vs *ARR7-llama GFPnes*, p = 1; *drmy1* vs *drmy1 ARR7-llama GFPnes*, p = 1.538×10^-7^). Data for *drmy1* were reused from Figure 2H, 2J. Data for *ARR7-llama GFP-nes* and *drmy1 ARR7-llama GFP-nes* were pooled from two independent lines (7-4 and 7-6). Sample size: WT, n = 78; *arr3,4,5,6,7,8,9,15*, n = 28; *ahp6*, n = 106; *ARR7-llama GFPnes*, n = 16; *drmy1*, n = 67; *drmy1 ARR7-llama GFP-nes*, n = 20. **(R)** Working model. In WT, DRMY1 maintains TOR activity and translation, which sustains the rapid production of cytokinin signaling inhibitors (ARR7 and AHP6) in response to cytokinin signaling. These inhibitors maintain cytokinin signaling at a normal level, allowing auxin and cytokinin signaling to interact and form robust spatial patterns. Robust patterning of auxin and cytokinin signaling gives rise to robustly initiated sepal primordia. In *drmy1*, due to decreased TOR signaling and translation rate, the meristem cannot rapidly produce cytokinin signaling inhibitor proteins in response to cytokinin signaling. As a result, cytokinin signaling is upregulated, which rescues the translation rate reduction in a homeostatic mechanism. This upregulation of cytokinin signaling disrupts the robust spatial pattern of both cytokinin and auxin signaling, which in turn causes variability in sepal initiation. Scale bars in all micrographs, 25 µm.

We started testing the level of protein reporters for a variety of cytokinin signaling components, in *drmy1* vs. WT and/or under translation or TOR inhibition. We reasoned that, given that the transcript level of most cytokinin signaling components do not significantly differ between *drmy1* and WT inflorescence tissue (Figure S7B), any changes in the level of these protein reporters should reflect post-transcriptional regulation, including mRNA translation. For A-type ARRs, we were unable to detect fluorescence in the inflorescence of a published *pARR4::ARR4-GFP* line^90^. We reasoned that this was because A-type ARRs have low protein levels (none was detected in our proteomics dataset) and short half-lives^91^. We therefore employed LlamaTagging, a recently developed method to visualize the abundance of nuclear-localized proteins with short half-lives^92^. Rapidly degraded proteins cannot be visualized through fusion with standard fluorescent proteins, because fluorescent proteins take time to mature before they fluoresce, and the protein of interest is degraded before the maturation of the fluorescent protein. On the other hand, the LlamaTag folds immediately. A LlamaTag with a high affinity for GFP can be encoded as a translational fusion with a nuclear-localized protein of interest. Soon after translation, the fusion immediately binds cytoplasm-localized GFP and translocates it to the nucleus. Thus, increased GFP fluorescence in the nucleus indicates higher abundance of the protein of interest (Figure 7B).

We focused on ARR7, the most highly expressed A-type ARR in our inflorescence RNA-seq (Figure S7B) which was also shown to be nuclear-localized^93,94^. We designed a construct with ARR7 fused with GFP-specific LlamaTag by a short linker, driven by the ARR7 native promoter (*pARR7::ARR7-linker-llama-ARR7ter*; *ARR7-llama* for short). This construct was co-transformed with cytoplasm-localized GFP containing a nuclear exclusion signal (*pUBQ10::sfGFP-nes-UBQ3ter*; *GFP-nes* for short; Figure 7C). As a proof of concept, we treated this reporter in WT background with 200 µM BAP. We found that GFP signal became more nuclear-localized within 5 hours of the treatment (Figure S7C-D), agreeing with an increased expression and stability of A-type ARR proteins upon cytokinin application as previously reported^91,95^.

We then compared the localization of GFP signal in floral meristems of WT and *drmy1* before sepal initiation. According to the null hypothesis, increased cytokinin signaling in *drmy1* would cause an increase in ARR7 protein level and thus more nuclear-localized GFP signal in *drmy1 ARR7-llama GFP-nes* than its WT counterpart. This is because cytokinin signaling increases the gene expression and protein stability of A-type ARRs^88,91,95,96^. In contrast, according to our hypothesis, insufficient protein synthesis of A-type ARRs contributes at least in part to increased in cytokinin signaling in *drmy1*, so we expect to see reduced ARR7 protein level and thus more cytoplasm-localized GFP signal in *drmy1 ARR7-llama GFP-nes*. We found that WT buds had slightly more nuclear-localized GFP signal than cytoplasm-localized GFP signal, with brighter spots corresponding to the nucleus surrounded by darker grooves in between corresponding to the cytoplasm (Figure 7D,E). In contrast, in the periphery of *drmy1* buds, GFP signal localizes more to the cytoplasm than to the nucleus, with darker spots surrounded by brighter grooves (Figure 7D,E). More nuclear GFP was present near the center of *drmy1* buds. This result indicates nuclear ARR7 protein concentration is reduced in the *drmy1* mutant, particularly in the zone where sepals initiate. To see whether this conclusion holds in other translation-inhibited conditions, we treated WT plants carrying the *ARR7-llama* and *GFP-nes* reporters with the translation inhibitor CHX and the TOR inhibitor AZD-8055. 2 µM CHX treatment for 24 hours drastically reduced the nuclear localization of the GFP signal and increased its cytoplasmic localization (Figure 7F). 2 µM AZD-8055 treatment for 72 hours had a milder but similar effect (Figure 7G). These treatments did not affect the localization of the GFP signal in plants without *ARR7-llama* (Figure S7E-H). These results show that conditions that decrease global protein synthesis greatly decrease the nuclear level of ARR7 protein.

We also tested whether TOR or translation inhibition alter the protein level of AHP6. To this end, we imaged the *pAHP6::AHP6-VENUS*^16^ protein reporter under mock, CHX, or AZD-8055 treatment. While mock-treated buds highly accumulate the AHP6 protein in the four incipient sepal primordia, buds treated with CHX or AZD-8055 abolished AHP6 accumulation within 72 hours (Figure 7H). The *pAHP6::GFP-ER*^97^ transcriptional reporter does not change under these treatments (Figure 7I), in agreement with our RNA-seq data of WT vs. *drmy1* (Figure S7B), suggesting that the change in AHP6 protein level is due to post-transcriptional regulation such as altered translation.

It is possible that inhibition of TOR and translation results in a general, uniform reduction in the level of all proteins, not just for the cytokinin signaling inhibitors ARR7 and AHP6, but also for the positive regulators of cytokinin signaling. For this, we tested whether AHP3, a component of the cytokinin phosphorelay^98,99^, respond to TOR and translation inhibition. CHX treatment did not affect the level or spatial localization pattern of the *pAHP3::AHP3-GFP* reporter^100^, while AZD-8055 treatment only mildly decreased its level (Figure S7I). We also tested how the level of other, more generic proteins respond to TOR and translation inhibition. Unexpectedly, the level of *pUBQ10::mCherry-RCI2A* increased upon 72 hours of CHX or AZD-8055 treatments (Figure S7J). Overall, these results show that TOR and translation inhibition does not result in a uniform reduction in the level of all proteins, but specific proteins such as ARR7 and AHP6 are dramatically decreased. Further, these results are consistent with our hypothesis that depletion of cytokinin signaling inhibitor proteins, including ARR7 and AHP6, may underlie the upregulation of cytokinin signaling when the floral meristem is under TOR or translation inhibition (Figure 4).

Having shown a reduction in the level of ARR7 and AHP6 greater than other proteins under TOR and translation inhibition, we next tested whether such a reduction contributes to the variability in sepal initiation also seen under such conditions (Figure 2). High-order mutant of A-type *ARR*s (*arr3,4,5,6,7,8,9,15*^101^) shows reduced size of the inner sepal primordium (Figure 7J,K) and a minor but significant increase in the positional variability of sepal primordia compared to WT (Figure 7Q), although sepal primordium number remains robust (Figure 7P). The *ahp6* mutant^16^ shows great variability of sepal primordium number, position, and size, although to a lower extent than *drmy1* (Figure 7J,L,P,Q). Similarly, if the reduction of ARR7 level in *drmy1* contributes to variability in sepal initiation, increasing ARR7 expression should restore sepal initiation robustness. While *ARR7-llama GFP-nes* plants phenocopy WT (Figure 7M), introducing the *ARR7-llama GFP-nes* constructs into *drmy1* plants partially restores robustness in sepal initiation, particularly in the position of sepal primordia (Figure 7N-Q). In older buds, while *drmy1* buds show great variability in sepal number and size resulting in gaps, some buds of *drmy1 ARR7-llama GFP-nes* have robustly sized sepals that are able to close properly (Figure S7K-N). Overall, these results show that reducing the level of cytokinin signaling inhibitor proteins ARR7 and AHP6 create variability in sepal primordium initiation, and increasing their level in *drmy1* partially restores robustness. We propose that, during hormone patterning prior to sepal initiation, the rapid synthesis of these inhibitor proteins in response to auxin and cytokinin signaling is crucial for maintaining the homeostasis of cytokinin signaling and thus the robustness in sepal initiation.

We also considered other hormone-related proteins that are dynamically regulated during organogenesis and thus may be depleted under translation defects. AUXIN/INDOLE-3-ACETIC ACID INDUCIBLE (Aux/IAA) proteins are auxin signaling inhibitors that are rapidly induced by auxin^102,103^. They bind auxin and are rapidly degraded by the ubiquitin E3 ligase SKP1, CUL1, F-BOX PROTEIN (SCF) complex involving TRANSPORT INHIBITOR RESPONSE1/AUXIN SIGNALING F-BOX (TIR1/AFB)^104–107^. Degradation is dependent on the Short Linear Motif (SLiM) degron contained within Domain II (DII)^106–108^. We hypothesized that the level of DII-containing proteins including Aux/IAAs would be drastically decreased in translation-inhibited conditions such as *drmy1* because they are unable to be rapidly synthesized to keep up with their degradation upon auxin signaling. To test this, we used the R2D2 reporter^109^, which contains a DII fused with 3xVENUS (*pUS7Y::DII-n3xVENUS*), and as a control, a mutated non-degraded DII fused with tdTomato (*pUS7Y::mDII-ntdTomato*). We compared this reporter in *drmy1* vs. WT. The ratio of *VENUS* to *tdTomato* was not reduced in *drmy1*, but instead slightly but significantly elevated (Figure S7O-P). In addition, *drmy1* has stochastic patches of DII-VENUS degradation, consistent with its often mislocalized auxin maxima (Figure 4A, S4B), unlike WT which had four patches of degradation corresponding to the four incipient sepal primordia where auxin maxima robustly form (Figure 4A, S4A). Overall, these results suggest that the level of DII-containing Aux/IAA proteins is not reduced in *drmy1*, despite the high requirement for synthesis due to their rapid turnover. They also indicate that not all proteins that are dynamically regulated in response to hormone signaling are equally affected by translation inhibition, which may result in different changes in hormone signaling output under such condition.

## DISCUSSION

Robustness, the strikingly reproducible development of phenotype, has fascinated biologists for decades^2^. The Arabidopsis flower robustly develops four sepals of equal size. This stems from the robust initiation of four sepal primordia from the floral meristem, which is in turn dictated by the robust patterning of auxin and cytokinin controlled by DRMY1^27^. Here we elucidated how DRMY1 controls robust hormone patterning and thus robust sepal initiation. We show that DRMY1 sustains TOR activity, ribosomal content, and translation. We further show that inhibition of TOR activity or translation is sufficient to cause variability in the timing, position, and number of sepal primordia, mimicking the *drmy1* phenotype. Our findings are in concert with previous studies that have shown robustness is often maintained by genes involved in central cellular processes^2^. In our case, the rate of translation in wild type maintains proper levels of ARR7 and AHP6, two cytokinin signaling inhibitor proteins, which need to be rapidly synthesized to dampen cytokinin signaling. Homeostasis of cytokinin signaling ensures robustness in auxin signaling patterns, and thus robustness in sepal initiation (Figure 7R, top). In the *drmy1* mutant, the reduced TOR activity, ribosomal content, and translation rate causes inability to rapidly synthesize these inhibitor proteins. Consequently, cytokinin signaling is elevated, disrupting the robust spatial pattern of auxin signaling, leading to variable sepal initiation (Figure 7R, bottom). Blocking cytokinin signaling in *drmy1* is sufficient to restore robust initiation of four sepal primordia, but has severe consequences on the overall fitness of the plant. Our results reveal how defects in hub cellular processes such as TOR signaling and translation can have tissue-specific phenotypic effects.

Although we propose that reduced TOR activity and mRNA translation affects developmental robustness through reduced synthesis of ARR7 and AHP6, we do not exclude other potential mechanisms that could contribute to the drmy1 phenotype. For example, we observed that several subunits of the 26S proteasome are more abundant in drmy1 than in WT (Supplemental Dataset 1), which could reflect or influence the accumulation of proteotoxic peptides and disrupt protein homeostasis^110–114^. This accumulation of 26S proteasomes could function upstream and/or downstream of the defect in TOR activity that impacts robust organogenesis^34,115–120^. Substantial future research will be needed to comprehensively define how DRMY1 participates in the complex interactions among TOR, mRNA translation, and proteolysis.

It was discovered long ago that extrinsic cytokinin application to plant tissue or cell-free extracts can promote mRNA translation^75–78^. Recent studies further confirmed that the up-regulation of translation by cytokinin is at least in part mediated by the cytokinin signaling pathway^74,82^. Here, we show that cytokinin signaling in floral buds is upregulated in translation-inhibited conditions, such as *drmy1*, AZD-8055 treatment, or CHX treatment (Figure 4), through reduced level of cytokinin inhibitor proteins (Figure 7; also potentially through other untested mechanisms). The enhanced cytokinin signaling maintains translation rate at a level necessary for the survival and reproduction of the plant (Figure 6). We propose that this represents a homeostasis mechanism where plants leverage increased cytokinin signaling to rescue the translation rate reduction caused by deficient TOR activity and ribosomal content (Figure 7R, bottom). It remains to be tested how widely this mechanism is applicable to other mutants with ribosomal defects, or whether parallel mechanisms operate in other species across kingdoms.

While translation-inhibited plants likely upregulate cytokinin signaling to maintain protein synthesis, this upregulation negatively affects developmental robustness. We have previously shown that exogenous cytokinin application to the WT floral meristem increases variability in PIN1 convergence and auxin signaling patterns, and consequently, in sepal initiation. These effects are more pronounced in the *drmy1* mutant, which by itself has increased and diffuse cytokinin signaling^27^. Here, we provide additional evidence that increased and diffuse cytokinin signaling is necessary for such variability. While *drmy1* and CHX-treated WT floral meristems are variable in auxin signaling pattern and sepal initiation (Figure 2, 4), mutations in *wol* and *arr1,10,12*, which decreases cytokinin signaling, largely restore robustness (Figure 5). Robustness is also restored in the mature sepals of *drmy1 wol* and *drmy1 arr1,10,12*, enabling sepal closure (Figure S6). Similar effects in restoring robustness are seen when an extra functional transgene of *ARR7* (*pARR7::ARR7-llama*) is introduced to the *drmy1* mutant (Figure 7J-Q, S7K-N). Our results suggest that cytokinin upregulation is necessary and sufficient for variability in auxin patterning and sepal initiation, indicating that the cytokinin signaling changes are primary defects in *drmy1*, and the auxin signaling changes are secondary. Our results suggest a mechanism different from that previously reported in *ul4y*, *ul18z*, and *el24y*, where ribosomal mutations affect auxin signaling through reduced translation reinitiation of uORF-containing mRNAs, including those of AUXIN RESPONSE FACTOR (ARF) 3, 5, and 7^64–66^. While we found that uORF-containing mRNAs generally have reduced protein-transcript ratio in *drmy1* suggestive of reduced translation, we did not see a consensus reduction in the level of uORF-containing promoter reporters of ARFs (Figure S5). This suggests that the variable auxin signaling pattern in *drmy1* is unlikely to result from changes in uORF-mediated translational regulation of ARFs. Overall, our results suggest that homeostasis in cytokinin signaling is crucial for maintaining robust patterns of auxin signaling and robust morphogenesis in the floral meristem.

Mutations affecting ribosome abundance or translation have long attracted interest due to the surprisingly tissue-specific phenotypes they cause^121^. In humans, these mutations have been associated with diseases collectively known as ribosomopathies, where patients show various abnormalities in blood, skeleton, hair, teeth, and pancreas, as well as intellectual disability and increased risk of cancer^122–127^. Ribosomal protein mutants have been characterized in numerous other species with similarly diverse impacts. They display a range of specific phenotypic changes, such as altered pigmentation and skeletal structure in mouse^128–130^ and zebrafish^131^, shorter bristles and notched wing margins in fruit fly^23,132^, abnormal gonad development in worm^133^, and pointed leaves and abnormal vascular patterning in Arabidopsis^64,66,134–136^. Here, we show that the Arabidopsis mutant *drmy1* has reduced TOR activity, ribosomal content, and translation rate, causing variable sepal initiation which phenocopies the ribosomal mutants *ul4y* and *ul18z* and the TORC1 assembly mutant *spaghetti-1* (Figure 2, 3). We therefore propose that *drmy1* is an Arabidopsis ribosomopathy mutant like those previously characterized^135^.

Several mechanisms have been proposed to explain why ribosomopathies do not usually cause a general reduction in growth, but rather affect development in tissue-specific ways. These include extra-ribosomal functions of certain ribosomal proteins^137–141^, altered translation behavior of ribosomal variants on certain mRNAs^142^, different competitiveness of mRNAs for scarce ribosomes^64–66,143–146^, and high translation rate requirement for certain proteins^147,148^. For example, neurotransmitter release in animals relies on constant synthesis of the synaptic vesicle protein Syt1^149^. A Drosophila *Minute* mutant, *uS15/+,* shows reduced synthesis of Syt1, which in turn reduces ecdysone secretion in 5-HT neurons, causing delayed larval-to-pupal transition^147^. Similarly, the human apoptosis inhibitor Mcl-1 has a half-life of ∼30 min and thus requires a high translation rate to maintain its proper level. Under translation inhibition, the synthesis of Mcl-1 is unable to keep up with its degradation, causing apoptosis^148^. Here, we show that the levels of ARR7 and AHP6, which are rapidly induced by cytokinin and auxin signaling, respectively^150,151^, are drastically reduced under translation inhibition, which underlies the upregulation of cytokinin signaling and loss of robustness in auxin signaling and morphogenesis (Figure 7R). This mechanism parallels those previously found in animal systems^147,148^, highlighting how downregulation of proteins with high translational requirements can underlie the tissue-specificity of ribosomopathy. Outside the floral meristem, the *drmy1* mutant shows other phenotypic changes such as enlarged shoot apical meristem, reduced apical dominance, phyllotaxy defects, and reduced root system, all of which are related to altered cytokinin/auxin signaling activity^27^. Thus, our work highlights how defects in translation, which occurs in every cell, can have tissue-specific effects on how cells robustly arrange into organs.

In addition, we note that not all proteins are equally reduced under broad translation inhibition. Our data suggest that the cytokinin signaling inhibitor proteins ARR7 and AHP6 are drastically reduced, compared to others such as AHP3, RCI2A, and GFP-nes. This suggests that the observed increase in cytokinin signaling under translation inhibition may be due to an imbalance in the relative levels of activators and inhibitors, which may further suggest that the inhibitor proteins are more temporally dynamic and thus have higher translational requirements during development. These hypotheses remain to be tested in future studies.

## ACKNOWLEDGEMENTS

We thank Bella Burda, Frances Clark, Byron Rusnak, Erich Schwarz, Avilash Yadav, and Maura Zimmermann for comments and suggestions on the manuscript. We thank Frank Wellmer for the *ap1 cal 35S::AP1-GR* (Ler) seeds, Elliot Meyerowitz and Arnavaz Garda for the *DR5::3xVENUS-N7*/*PIN1::GFP* (Ler) seeds, Teva Vernoux and Géraldine Brunoud for the *TCS::GFP*, *TCS-DR5* (Col), *ahp6*, *pAHP6::AHP6-VENUS*, and *pAHP6::GFP-ER* seeds, Jan Smalle for the *arr1-1 35S::ARR1* seeds, Joseph Kieber and Jamie Winshell for the *arr3,4,5,6,7,8,9,15* and *pARR4::ARR4-GFP* seeds, Thomas Greb and Min-Hao Chiang for the *pARF5::ER-ARF5-HDEL* seeds, and Jianru Zuo for the *pAHP3::AHP3-GFP* plasmid. We thank Georg Jander for advice and protocol for cytokinin extraction. We thank Brian Curtis and Frank Schroeder for help on mass spectrometry, and Sheng Zhang and Qin Fu for help on Xcalibur. We thank Richie Ragas, Yanã Rizzieri, and Ziqing Wei for assistance on experiments and data analysis. We thank Vicky Spencer and Minsung Kim for sharing the *in vivo* Torin2 treatment protocol. We thank Arabidopsis Biological Resource Center for providing seed stocks and plasmids used in this research.

Research reported in this publication was supported by the National Institute of General Medical Sciences of the National Institutes of Health (NIH) under award numbers R01GM134037, DP5OD023072, and R01GM145814; Cornell Graduate School new student fellowship (S.K.); and in part by a Schmittau-Novak Grant from the School of Integrative Plant Science, Cornell University (M.Z.). H.G.G. was supported by NIH Director’s New Innovator Award (DP2 OD024541-01) and NSF CAREER Award (1652236), NIH R01 Award (R01GM139913), and the Koret-UC Berkeley-Tel Aviv University Initiative in Computational Biology and Bioinformatics. H.G.G. is also a Chan Zuckerberg Biohub Investigator. We thank the Metabolomics Workbench^152^ for providing the data repository for our cytokinin mass spectrometry data, which is funded by NIH under award numbers U2C-DK119886 and OT2-OD030544. We thank the Biotechnology Resource Center (BRC) Genomics Facility (RRID:SCR_021727) of Cornell University for performing RNA-seq on *ap1 cal AP1-GR* inflorescence samples. We thank Novogene for performing seedling RNA-Seq library synthesis and sequencing. We thank the BRC Proteomics Facility (RRID:SCR_021743) of Cornell University for performing mass spectrometry on *ap1 cal AP1-GR* inflorescence samples, and NIH SIG Grant 1S10 OD017992-01 for supporting the Orbitrap Fusion mass spectrometer. The content is solely the responsibility of the authors and does not necessarily represent the official views of the National Institutes of Health.

## AUTHOR CONTRIBUTIONS

S.K.: Conceptualization, Investigation, Formal Analysis, Visualization, Writing - Original Draft, Writing - Review & Editing. M.Z.: Conceptualization, Investigation, Formal Analysis, Writing - Review & Editing. M.R.S: Investigation, Visualization. D.P.: Investigation, Formal Analysis, Visualization. L.J.: Investigation. R.E.M.: Investigation, Formal Analysis. S.A.: Methodology, Resources. B.V.L.V.: Methodology, Resources, Writing - Review & Editing. H.G.: Methodology, Resources, Writing - Review & Editing, Supervision. S.B.Q: Writing - Review & Editing, Supervision. J.O.B.: Resources, Writing - Review & Editing, Supervision. A.H.K.R: Conceptualization, Writing - Review & Editing, Supervision, Project administration, Funding acquisition.

## MATERIALS AND METHODS

### Plant material

Most Arabidopsis plants were in Col-0 background (WT). *ap1 cal 35S::AP1-GR* was in Ler background. *drmy1* (Col-0) was backcrossed to Ler twice and then crossed with *ap1 cal 35S::AP1-GR* to obtain *drmy1 ap1 cal 35S::AP1-GR*. R2D2 was originally in Col-Utrecht background and was backcrossed twice into WT (Col-0) and *drmy1* (Col-0). The following mutants and reporters were previously described: *drmy1-2*^27^, *wol-1*^153^, *spaghetti-1* (*tpr5-1*)^154^, *ahp6*^16^, *arr3,4,5,6,7,8,9,15*^101^, *ap1 cal 35S::AP1-GR* (Ler)^35,36^, *arr1-1 35S::ARR1*^74^, *DR5::3xVENUS-N7*^155^, *TCS::GFP*^156^, *pARF5::ER-EYFP-HDEL*^157^, *pUS7Y-mDII-NtdTomato-pUS7Y*-*DII-N3xVENUS* (R2D2)^109^, *35S::mCirtine-RCI2A*^27^, *UBQ10::mCherry-RCI2A*^27^, *pAHP3::AHP3-GFP*^100^, *pAHP6::AHP6-VENUS*^16^, and *pAHP6::GFP-ER*^16,97^. The following mutants and reporter lines were obtained from Arabidopsis Biological Resource Center (ABRC): *ul4z* (SALK_130595), *ul4y* (SALK_029203), *ul18z* (SALK_089798), *arr1-3 arr10-5 arr12-1*^158^ (CS39992), *lst8-1-1* (SALK_002459), *pARF3::N3xGFP*^159^ (CS67072), *pARF6::N3xGFP*^159^ (CS67078), *pARF8::N3xGFP*^159^ (CS67082), *pARF10::N3xGFP*^159^ (CS67086).

### Llama-tagged ARR7 construct

For the LlamaTag system, we first generated plasmid *pVV13* containing linker-llama. We amplified the LlamaTag (from a plasmid containing vhhGFP4^160^) and added a linker sequence of tccggagcagctgcggctgccgctgcggcagcggccactagt at its 5’ end by two rounds of overlap PCRs. Primers for the first round were oVV64 and oVV53, and primers for the second round were oVV35 and oVV53. After the second round, we A-tailed the PCR product according to the Promega manufacturer’s protocol. A-tailed product was ligated to the pGEMTeasy vector according to the Promega ligation protocol, to create the plasmid *pVV13*.

To make *pARR7::ARR7-llama*, a genomic fragment of *pARR7::ARR7* minus the stop codon and terminator was amplified from the Arabidopsis (Col-0) genome using the primers oSK197 and oSK198. The linker-llama fragment was PCR-amplified from *pVV13* using the primers oSK199 and oSK200. The *ARR7* stop codon, 3’ UTR, and terminator was amplified from the Arabidopsis (Col-0) genome using the primers oSK201 and oSK202. *pMLBART* backbone was digested with NotI, and all fragments were assembled into *pMLBART* using NEBuilder according to the manufacturer’s protocol.

To make *pUBQ10::sfGFP-NES:UBQ3ter*, sfGFP sequence was amplified from the *35S-sfGFP-nosT* plasmid^161^ (Addgene # 80129) using primers UsfGM-F1 and UsfGnes-R1. The UBQ10 promoter was amplified from the *UPG* plasmid^162^ (Addgene # 161003) using primers OutALFd and UsfGM-R1. The UBQ3 terminator was amplified from the *UPG* plasmid^162^ (Addgene # 161003) using primers UsfGnes-F1 and OutALRb. Primer overhangs spanning the junction between sfGFP and the UBQ3 terminator contain the sequence of the mouse PKIα NES. *pCambia1300* backbone was digested with BamHI and KpnI, and all fragments were Gibson-assembled into the backbone. Sequences of primers, *pARR7::ARR7-llama*, and *pUBQ10::sfGFP-NES:UBQ3ter* can be found in Supplemental Dataset 5.

Col-0 plants were co-transformed with *pARR7::ARR7-llama* and *pUBQ10::sfGFP-NES:UBQ3ter*, and selected with Basta (for *pARR7::ARR7-llama*) + Hygromycin (for *pUBQ10::sfGFP-NES:UBQ3ter*). Surviving T1 plants were screened for clear nuclear signal in the inflorescence, and 5 independent T1 plants were selected and crossed into *drmy1*. F2 plants from each line were again selected with Basta + Hygromycin and genotyped. One line showed co-segregation with the *DRMY1* locus. Two lines showed severe silencing in the F2 and could not be used. Two lines (7-4 and 7-6), though with minor silencing in F2, were used for imaging and image analysis. F3 plants of 7-4 and 7-6 had severe silencing, and therefore only F2 were imaged.

### Plant growth conditions

For most experiments, seeds were sown in wetted Lamber Mix LM-111 soil and stratified at 4°C for 3-5 days. For experiments including *drmy1 wol* and *drmy1 arr1,10,12*, all seeds were sown onto ½ MS plates with 0.05% (w/v) MES, 1% (w/v) sucrose, 1.2% (w/v) agar, pH 5.7, and stratified at 4°C for a week. They were grown for 7-10 days before being transplanted to soil (for imaging of inflorescence or aerial part of the plant) or left on the plates until desired time of the experiment (for seedling imaging or puromycin labeling).

Most plants were grown under 16 h – 8 h light-dark cycles (fluorescent light, ∼100 µmol m^-1^ s^-1^) at 22°C in a Percival walk-in growth chamber. We found that the *drmy1* phenotype is more pronounced in this condition than under continuous light. The *ap1 cal 35S::AP1-GR* and *drmy1 ap1 cal 35S::AP1-GR* plants were grown in soil under continuous light at 16°C to prevent premature floral induction.

### Flower staging

Flower buds were staged as previously described^37^. Briefly, stage 1 is when the floral meristem emerges, but not yet separated, from the inflorescence meristem. Stage 2 is when the floral meristem separates from the inflorescence meristem but with no floral organs initiated. Stage 3 is when sepal primordia initiate. Stage 4 is when sepal primordia bend to cover part of the floral meristem. Stage 5 is when stamen primordia initiate. Stage 6 is when sepal primordia completely cover the floral meristem.

### RNA-seq data collection and analysis

For RNA-seq in the inflorescence, bolting *ap1 cal 35S::AP1-GR* and *drmy1 ap1 cal 35S::AP1-GR* plants were induced daily with an aqueous solution containing 10 µM dexamethasone (Sigma-Aldrich), 0.01% (v/v) ethanol, and 0.015% (v/v) Silwet L-77 (Rosecare.com). When sepals initiated from the floral meristems, usually on the fourth day after three daily inductions, three inflorescence samples per genotype (including inflorescence meristems and buds under stage 6) were collected and immediately put into liquid nitrogen. RNA extraction, library preparation, RNA-seq, and data analysis for inflorescence samples were done as previously described^27^ with a few changes. After read mapping, genes with at least two raw reads in at least two biological replicates in either WT or *drmy1* were kept for downstream analysis. For differentially expressed genes, we set a log2 fold change threshold of ±1 and a BH-adjusted p-value threshold of 0.05. For GO term enrichment, gene-GO mapping data was obtained from TAIR (https://www.arabidopsis.org/download_files/GO_and_PO_Annotations/Gene_Ontology_Annotations/ATH_GO_GOSLIM.txt). The R package “topGO”^163^ (version 2.38.1) was used for the enrichment, with statistic “fisher”, algorithm “weight01”, annotation function “annFUN.gene2GO”, and minimum node size 10. The results were ranked by their p-value, and the first 8 terms were plotted.

For RNA-seq in seedlings, WT and *drmy1* seedlings were grown to quiescence (7 days) in ½ MS liquid media as previously described^34^. After 7 days, the media was replaced with ½ MS liquid media containing 15 mM glucose and incubated for 24 hours to activate TOR. Seedlings were then incubated with or without AZD-8055 in addition to 15 mM glucose in ½ MS liquid media for 2 hours before collecting tissue. RNA was extracted from 100 mg pooled seedlings using the Spectrum Plant Total RNA Kit (Sigma). This RNA was used as a template for RNA-Seq library synthesis and sequencing, which was performed by Novogene. RNA-seq data for AZD-8055 treated WT and *drmy1* seedlings were preprocessed with fastp (v. 0.22.0) using default parameters. Preprocessed reads were then mapped to the TAIR10 reference genome using STAR (v. 2.7.10z_alpha_220314). Following alignment, BAM output files from STAR were used to generate feature counts for transcripts using subread-featureCounts (v. 2.0.3) and the Araport11 transcriptome. TPMs were generated using TPMCalculator (v. 0.0.3). Differential expression analysis was performed using feature count data and DESeq2 (v. 1.36.0).

A list of genes with uORFs based on gene models of the TAIR10 Arabidopsis genome assembly were downloaded from von Arnim et al.^69^. For each gene, within each genotype, protein-transcript ratio was calculated as the ratio between mean protein abundance and mean transcript TPM across all bio-reps in our proteomics and RNA-seq datasets, respectively. This was log2-transformed, and the difference between *drmy1* and WT was calculated. This was used as an indicator of translation rate difference between *drmy1* and WT, although we acknowledge that other factors such as protein stability may affect this number. This was plotted against the number of uORFs in each gene model (0, 1, or ≥ 2).

### Proteomics

Five induced inflorescence samples of WT and *drmy1* in *ap1 cal AP1-GR* background were collected as described above. Samples were ground in liquid nitrogen. Total soluble proteins were extracted in ice-cold extraction buffer (50 mM PBS-HCl (pH 8.0) buffer with 150 mM NaCl, 2% NP-40, 1 mM PMSF, 1x Roche cOmplete protease inhibitor cocktail (Sigma 11697498001), and 1x Halt TM Phosphatase inhibitor cocktail (ThermoFisher 78420)) and filtered through Pierce™ Micro-Spin Columns (30 µm pore size; Thermo Scientific 89879). Extracts were RuBisCO-depleted using Seppro Bubisco Kit (Sigma SEP070-1KT), concentrated, denatured, reduced, cysteine blocked, trypsin-digested, and TMT 10-plex labeled. Then, mass spectrometry was done using an UltiMate 3000 RSLCnano / Orbitrap Fusion system (Thermo Scientific). Raw data was searched against the NCBI protein database using PD 2.3 (Thermo Scientific) with Sequest HT searching engine. Precursor-based protein identification and relative quantification was done using the standard processing workflow in PD 2.3, with an additional node of Minora Feature Detector. Proteins with at least 2 supporting peptides were kept for downstream analysis. For each protein, data was fit with an ANOVA model and a p-value was calculated. Proteins with a p-value < 0.05 were considered differentially accumulated in *drmy1*. GO term enrichment was done as above, using genes corresponding to the differentially accumulated proteins.

### Polysome extraction and profiling

Three induced inflorescence samples of WT and *drmy1* in *ap1 cal AP1-GR* background were collected as described above, and polysomes were extracted as previously described^164^. Briefly, samples were ground in liquid nitrogen, mixed with an extraction buffer twice the volume of pulverized tissue (0.2 M Tris pH 9.0, 0.2 M KCl, 0.025 M EGTA, 0.035 M MgCl_2_, 1% (w/v) Brij-35, 1% (v/v) Triton X-100, 1% (v/v) Igepal CA-630, 1% (v/v) Tween-20, 1% (w/v) Sodium deoxycholate, 1% (v/v) Polyoxyethylene 10 tridecyl ether, 5 mM Dithiothreitol, 1 mM Phenylmethylsulfonyl fluoride, 100 µg/ml cycloheximide, 100 µg/ml chloramphenicol, 40 U/ml RNasin, 10 U/ml DNase I), and let sit on ice for 10 min. Samples were centrifuged at 4°C 4,000 g for 5 min, supernatant was transferred to a new tube, centrifuged at 4°C 16,000 g for 15 min, and supernatant was filtered through Miracloth.

Polysome extracts were profiled as previously described^165^. Briefly, 600 µl of each sample were loaded onto a 15%-45% sucrose density gradient and centrifuged at 4°C 32,000 rpm in a SW41 rotor. Separated samples were fractionated at a rate of 0.375 mL/min in an Isco fractionation system, and absorbance at 254 nm was recorded.

### Puromycin labeling

Puromycin labeling was done as previously described^39^, with slight modifications.

In seedlings, when comparing WT and *drmy1*, in order to control for plant size, WT seedlings were grown for 8 days and *drmy1* seedlings were grown for 10 days (Figure 1E). When comparing WT, *drmy1*, *wol*, and *drmy1 wol*, we were unable to control for plant size because *drmy1 wol* seedlings were too small. We therefore controlled for plant age, and seedlings were grown to specified age (8 days for Figure 6B and 14 days for Figure 6C). Seedlings were harvested from plates and incubated with an incubation buffer (½ MS, 0.05% (w/v) MES, 1% (w/v) sucrose, 0.1% (v/v) Tween-20, 0.1% (v/v) DMSO, 1x Gamborg vitamin mix, pH 5.7), with or without 50 µM CHX, for 4 hours in an illuminated growth chamber. Then, the buffer was replaced with a fresh incubation buffer (which is same as above, but contains 50 µM puromycin (GoldBio P-600-100)), and incubation continued for another 45 min.

In inflorescences of WT and *drmy1* in *ap1 cal AP1-GR* background, inflorescences were DEX-induced as described above. Inflorescence samples were collected and put in an incubation buffer (½ MS, 1% (w/v) sucrose, 0.02% (v/v) Silwet L-77, 0.1% (v/v) DMSO, 50 µM puromycin, 1x Gamborg vitamin mix, pH 5.7), with or without 100 µM CHX. Samples were vacuum infiltrated for 15 minutes and then put on a rocking shaker in an illuminated growth chamber for 45 minutes. In both cases, at the end of the incubation, samples were washed three times with water, blot dry, weighed, and frozen in liquid nitrogen. Soluble proteins were extracted as described above. Puromycin incorporated into the proteins were detected in a Western blot using a mouse-origin anti-puromycin monoclonal antibody (12D10, Sigma MABE343, lot # 3484967) and a goat-anti-mouse HRP-conjugated secondary antibody (Abcam ab6789, lot # 3436981). RuBisCO large subunit in Ponceau S-stained membrane was used as a loading control. Quantification was done in ImageJ. A background signal was determined using blank regions, and subtracted from all quantified signals (separately for puromycin and Ponceau S).

### TOR activity assay

WT and *drmy1* seedlings were grown in a six-well plate containing ½ MS liquid media. After seven days, the media were replaced with half-strength MS liquid media plus 15 mM glucose and incubated for 24 hours. At least 120 quiescent seedlings per sample were collected and frozen in liquid nitrogen. Protein was then extracted from the plant tissue in 100 mM MOPS (pH 7.6), 100 mM NaCl, 5% SDS, 0.5% b-mercaptoethanol, 10% glycerin, 2 mM PMSF, and 1x PhosSTOP phosphatase inhibitor (Sigma). S6K-pT449 was detected by Western blot using a phosphospecific antibody (Abcam ab207399) and an HRP-conjugated goat anti-rabbit IgG secondary antibody (Jackson Immuno Research 111-035-003). Total S6K was detected using a custom monoclonal antibody described by Busche et al.^166^. Total protein visualized in Ponceau S-stained membrane was used as a loading control.

### Confocal microscopy

Confocal imaging of reporter lines in the inflorescence were done as previously described^27^. Briefly, main inflorescences (not side branches) were cut and dissected with a Dumont tweezer (Electron Microscopy Sciences, style 5, no. 72701-D) to remove buds older than stage 9 or 10. The inflorescences were then inserted upright into a small petri dish (VWR, 60 x 15 mm) containing inflorescence culture medium (1/2 MS, 1% (w/v) sucrose, 1x Gamborg vitamin mixture, 0.1% (v/v) plant preservative mixture (Plant Cell Technology) 1% (w/v) agarose, pH 5.8), leaving most of the stem inside the medium and the buds outside. They were then further dissected to reveal stage 6 and younger buds, immersed with water, and imaged under a Zeiss710 upright confocal microscope with a 20x Plan-Apochromat water-dipping lens (1.0 NA). For live imaging experiments, inflorescence samples were put in a continuous-light growth chamber between time points. To prevent bacterial growth, samples were transferred onto fresh media every 2 to 3 days, and for live imaging experiments lasting longer than 6 days, once in the middle, plants were incubated with an aqueous solution of 100 µg/ml Carbenicillin (GoldBio, C-103-5, lot # 0129.091814A) for 30 minutes.

To visualize tissue morphology of inflorescence samples without a reporter, samples were stained for 5 minutes with an aqueous solution of 0.1 mg/ml propidium iodide (PI) and 0.1% (v/v) Tween-20, washed three times with water, and imaged.

The following laser and wavelength were used in confocal imaging. Chlorophyll, excitation 488 nm, emission 647-721 nm. PI, excitation 514 nm, emission 566-659 nm. mCherry, excitation 594 nm, emission 600-659 nm. tdTomato, excitation 561 nm, emission 566-595 nm. For EYFP/VENUS/mCitrine, in *35S::mCirtine-RCI2A*, excitation 514 nm, emission 519-580 nm; in *DR5::3xVENUS-N7*, excitation 514 nm, emission 519-569 nm; in *pARF5::ER-EYFP-HDEL*, excitation 514 nm, emission 519-550 nm; in *R2D2*, excitation 488 nm, emission 493-551 nm. For GFP/sfGFP, in *pARR7::ARR7-llama UBQ10::sfGFP-NES*, excitation 488 nm, emission 493-569 nm; in *pARF3::N3xGFP*, *pARF6::*N3xGFP, *pARF8::N3xGFP*, and *pARF10::N3xGFP*, excitation 488 nm, emission 493-564 nm; in *TCS::GFP*, excitation 488 nm, emission 493-513 nm.

### Visualization of tissue morphology

For single-channel image stacks intended for the visualization of tissue morphology (*35S::mCitrine-RCI2A* or PI), stacks were 3D-rendered using the ZEN confocal software (Processing -> 3D). Parameters were set to best visualize tissue morphology, typically, minimum 5-10, ramp 60-80, maximum 100. Buds were rotated to desired orientation, and screenshots were taken using the “Create Image” button. For fluorophores that are dimmer, less sharp, or have a noisy background (*UBQ10::mCherry-RCI2A* or Chlorophyll), stacks were converted from LSM to TIF using ImageJ^167,168^, loaded into MorphoGraphX^169^, and screenshots were taken using the built-in screenshot function in MorphoGraphX.

To aid visualizing tissue morphology and determine the timing of sepal initiation, each stack was fitted with a surface, and a Gaussian curvature heatmap was calculated from the surface (see below). We consider a sepal primordium as initiated when we see a dark red band of positive Gaussian curvature (primordium) separated from the center of the floral meristem by a dark blue band of negative Gaussian curvature (boundary)^27^.

Gaussian curvature heatmaps were calculated as previously described^27^, with slight modifications. Briefly, stacks underwent the following processes in MorphoGraphX: Gaussian blur (3 times; X/Y/Z sigma = 1 µm for the first 2 times, and 2 µm for the third time), edge detection (threshold = 2000-8000 depending on the brightness of the stack, multiplier = 2.0, adapt factor = 0.3, fill value = 30000), marching cube surface (cube size = 8 µm, threshold = 20000), subdivide mesh, smooth mesh (passes = 5), subdivide mesh, smooth mesh (passes = 5), project mesh curvature (type = Gaussian, neighborhood = 10 µm, autoscale = no, min curv = -0.0015, max curv = 0.0015). For ease of visualization, the lookup table “jet” was applied to the mesh.

### Quantification of sepal initiation robustness

For sepal primordium number, screenshots were taken of stage 3-6 buds of indicated genotypes, in either ZEN or MorphoGraphX. The number of sepal primordia initiated were counted from these screenshots.

For variability in sepal primordium positioning, within each bud, an angular distance was measured between each pair of adjacent sepal primordia (with vertex at the center of the bud), using ImageJ. Note that the last pair was not measured – the angular distance was calculated as the sum of all other angular distances subtracted from 360°. A CV value (standard deviation divided by mean) was calculated from all the measured or calculated angular distances. Buds with sepal primordia evenly distributed around the bud periphery should have a small CV value, i.e. all angles are around 90° for four-sepal buds (or 72° for five-sepal buds, etc.). Buds whose sepal primordia distributed variably or randomly around the bud periphery will have widely varying angular distances between adjacent sepal primordia, and thus large CV values.

Relative sepal initiation timing was quantified as previously described^27^. Briefly, dissected inflorescence samples were live-imaged every 6 hours. A Gaussian curvature heatmap was generated for each sample at each time point and was used to determine the time point at which a sepal primordium initiates. A sepal primordium is considered initiated at time point Tn if it is absent at time point T(n-1) but becomes present at time point Tn. Within the same bud, we counted the number of time points between outer and inner sepal initiation, and between outer and lateral sepal initiation, and multiplied them by 6 hours to get the relative initiation timing of these sepals.

### Quantification of fluorescent reporters

For *TCS::GFP*, *pARF3::N3xGFP*, *pARF5::ER-YFP-HDEL*, *pARF6::N3xGFP*, *pARF8::N3xGFP*, *pARF10::N3xGFP*, *pUS7Y::mDII-NtdTomato, pUS7Y::DII-N3xVENUS*, and *UBQ10::mCherry-RCI2A*, total signal (integrated density) was quantified from maximum intensity projection images using ImageJ^167,168^. Fluorescence intensity was measured in pixel intensity units (0-255 range). Signal intensity was calculated as total signal divided by area.

For both *TCS::GFP* and *DR5:3xVENUS-N7*, circular histogram analysis was done as previously described^27^. Briefly, individual buds were cropped out of image stacks, channels were split using FIJI and saved in TIF format, and TIF stacks were imported into MorphoGraphX. Signal from outside the buds (e.g. inflorescence meristem, parts of other buds within the same image) was manually removed using the Voxel Edit function. Buds were positioned so that the incipient sepal primordia are in the XY plane: the incipient outer sepal is at 45°, the incipient inner sepal and the inflorescence meristem are at 225°, and the incipient lateral sepals are at 135° and 315°, respectively. Fluorescence intensity was measured in pixel intensity units (0-255 range). A circular histogram of bin width 1° centered around the Z axis was exported for each replicate expressing DR5 and/or TCS. Multiple circular histograms of the same reporter and genotype were pooled and mean ± SD were plotted.

For GFP signal in plants carrying *pUBQ10::sfGFP-nes-UBQ3ter* and *pARR7::ARR7-linker-llama-ARR7ter* reporters, screenshots were taken in MorphoGraphX as described above. Screenshots were subtracted of a background determined using blank regions with no tissue, and brightened to the same level to reveal differences in GFP distribution patterns. A square region containing 5-10 cells were taken from each screenshot, and GFP intensity (in gray value ranging from 0 to 255) along a straight line of 239 pixels in length was quantified using ImageJ^167,168^ (Analyze -> Plot profile). For ease of visualization, the curves were smoothed by taking the average of the gray value of 11 neighboring pixels (including itself) as the value of each pixel.

For VENUS or GFP signal in *pAHP6::AHP6-VENUS*, *pAHP6::GFP-ER*, *pAHP3::AHP3-GFP*, and *pUBQ10::mCherry-RCI2A* under mock, CHX, or AZD-8055 treatment, total signal at 24 hours (for CHX) or 72 hours (AZD-8055) was normalized by bud area in the 2D projection to get the signal intensity. To account for bud-to-bud differences in signal intensity prior to treatment, the signal intensity was normalized to the 0-hour time point (pre-treatment). Relative level between treatment and mock was calculated by normalizing this value to the mean of mock.

### *In vitro* drug treatments on inflorescence samples

For cycloheximide (CHX) treatment, a stock solution of 10 mM CHX was made from powder (Sigma C1988) in pH 4.0 water. The stock was filter-sterilized and stored in -20°C, and added to autoclaved inflorescence culture medium to a final concentration of 2 µM just before use. For AZD-8055 treatment, a stock solution of 16 mM AZD-8055 was prepared from powder (Cayman Chemical 16978) in DMSO within days of use, and stored in -80°C. The stock was added to autoclaved inflorescence culture medium to a final concentration of 2 µM. 0.0125% (v/v) DMSO was added to the mock medium. For 6-benzylaminopurine (BAP) treatment, a stock solution of 50 mM BAP was prepared from powder (Alfa Aesar A14678) in DMSO, and stored in -80°C. The stock was added to autoclaved inflorescence culture medium to a final concentration of 5 µM. 0.01% (v/v) DMSO was added to the mock medium.

Inflorescences were dissected and inserted into regular inflorescence culture medium without drugs, and pre-treatment image stacks were captured. Then, they were transferred into specified treatment or mock media, and imaged at the specified time points. For live imaging, inflorescence samples were transferred onto new medium after each imaging session.

### *In vivo* Torin2 treatment

Starting at 14 days after germination, twice each day for 15 days, 2 nmol of Torin2 (Cayman Chemical 14185) in 20 µl of aqueous solution containing 0.5% DMSO and 0.5% Tween-20 was applied to the center of the rosette using a pipette. For mock, 20 µl aqueous solution containing 0.5% DMSO and 0.5% Tween-20 was applied. At the end of the 15-day treatment period, inflorescences were dissected and put in the inflorescence culture medium for imaging.

To prevent Torin2 degradation, throughout the duration of this experiment, the Torin2 stock solution in DMSO was kept in -80°C and replaced each week, and the treatment and mock solutions were kept in 4°C and replaced each day.

### Imaging of whole plant, whole inflorescence, silique, and mature sepals

For whole-plant imaging, aerial parts of the plants were removed from the pots, flattened, put on a dark cloth, and imaged with a cell phone (iPhone 12, iOS 16.2).

For whole-inflorescence imaging, inflorescences consisting of open flowers and unopened buds were removed from the plant and held with forceps. Images were taken under a Zeiss Stemi 2000-C Stereo Microscope with a cell phone (iPhone 12, iOS 16.2).

For silique imaging, siliques on inflorescences sufficiently distant from the shoot apex that were developed and started to ripen were picked with forceps, opened with a razor blade, and imaged under a Zeiss Stemi 2000-C Stereo Microscope with a cell phone (iPhone 12, iOS 16.2).

Mature sepal imaging was done as previously described^26,27^. Briefly, mature sepals from stage 15 flowers (10^th^ to 25^th^ flower on the inflorescence) were dissected and sandwiched between two slides to flatten. Images were taken using a Canon Powershot A640 camera attached to a Zeiss Stemi 2000-C Stereo Microscope. Minor damages were manually fixed, and undesired objects such as pollen grains were manually removed from these images. Sepals with major damages were discarded. Then, a contour was extracted from each sepal using custom python scripts^26^. This gave us measurements such as length, width, area, etc. of each sepal. To measure between-flower variability of length, within each genotype and for each of outer, inner, and lateral positions, a CV (standard deviation divided by mean) of all sepals was calculated (for example, a CV of length of all outer sepals in WT). To determine statistical significance, genotypes were compared pairwise using permutation tests. To measure within-flower variability of length, a CV was calculated for all sepals within each flower (for example, a CV of length of outer, inner, and two lateral sepals in WT bud #10). For accurate calculation of CV, flowers with length data of at least four sepals were included in the analysis. To determine statistical significance, genotypes were compared pairwise using Wilcoxon rank sum tests.

### Cytokinin extraction and measurement

Cytokinin extraction was based on a previously published protocol^170^ with modifications. Briefly, five inflorescence samples of induced *ap1 cal 35S::AP1-GR*, and six inflorescence samples of induced *drmy1 ap1 cal 35S::AP1-GR* were collected as described above. Samples were ground in liquid nitrogen and twice extracted in methanol : water : formic acid (15:4:1). 200 pg of BAP per sample was added as an internal control. Extracts were centrifuged at 14,650 rpm in -4°C for 30 min, and supernatant was evaporated of methanol and reconstituted in 1% (v/v) acetic acid. Samples were passed through an Oasis MCX SPE column (Waters 186000252), washed with 1% acetic acid, washed with methanol, and eluted with 0.35 M ammonia in 70% methanol. Eluents were evaporated to complete dryness, reconstituted in 5% acetonitrile, and sent for LC-MS.

LC-MS was done as previously described^171^, with modifications. Briefly, 1 µl of each sample was injected into a Thermo Fisher Scientific Vanquish Horizon UHPLC System coupled with a Thermo Q Exactive HF hybrid quadropole-orbitrap high-resolution mass spectrometer equipped with a HESI ion source. Samples were separated on a C18 ODS column (AQUITY UPLC BEH C18, 1.7 μm, 2.1 × 100 mm, Waters), at a flow rate of 0.3 ml/min, with linear gradients of solvent A (0.1% formic acid) and solvent B (0.1% formic acid in methanol) according to the following profile: 0 min, 99.0% A + 1.0% B; 4.0 min, 55.0% A + 45.0% B; 7 min, 30.0% A + 70.0% B; and then with isocratic conditions: 8 min, 1.0% A + 99.0% B; 12 min, 99.0% A + 1.0% B. Cytokinins were detected using the positive ion mode.

For tZ, tZR, iP, iPR, and the internal control BAP, peaks were identified from an external standard mix composed of 0.1 µg/ml each of BAP (Alfa Aesar A14678), tZ (Sigma Z0876), tZR (Sigma Z3541), iP (Cayman Chemical 17906), and iPR (Cayman chemical 20522) in 5% acetonitrile. For cZ and cZR, peaks were identified based on previously reported precursor m/z and retention time^172^. Using Xcalibur (Thermo Scientific), peak area was quantified for each cytokinin in each sample, normalized against the peak area of BAP (internal control) and sample fresh weight, and then normalized against the average abundance of tZ in WT samples.

### Software

Image processing was done in ImageJ (version 2.9.0/1.53t, build a33148d777)^167,168^ and MorphoGraphX (version 2.0, revision 1-294, CUDA version 11.40)^169^.

Data processing was done in RStudio (R version 4.0.5 “Shake and Throw” (2021-03-31))^173^. Graphs were made using the package ggplot2 (version 3.3.3)^174^. Fisher’s contingency table tests were done using the function fisher.test in R. Wilcoxon rank sum tests were done using the function wilcox.test in R. Hypergeometric tests were done using the function phyper in R. Data fitting with ANOVA was done using the function aov in R. Figures were assembled in Adobe Illustrator (version 25.4.1). An RGB color profile “Image P3” was used for all the figures.

### Accession numbers

RNA-seq data for *ap1 cal AP1-GR* and *drmy1 ap1 cal AP1-GR* inflorescence tissue were deposited in Gene Expression Omnibus (GEO) under accession number GSE230100. RNA-seq data for WT and *drmy1* seedlings treated with mock or AZD-8055 were deposited in NIH BioProject under accession number PRJNA961813. Mass spectrometry data for proteomics were deposited in the ProteomeXchange Consortium via the PRIDE^175^ partner repository under accession number PXD041723 (reviewer username, and password: 8pl3ZD1l). Mass spectrometry data for cytokinins were deposited in NIH’s National Metabolomics Data Repository (NMDR) website, the Metabolomics Workbench^152^, under accession number ST002571.

## SUPPLEMENTAL INFORMATION TITLES AND LEGENDS

**Supplemental Figure 1. Evidence that the *drmy1* mutant has ribosomal and translation defects, associated with Figure 1.**

**(A)** The *drmy1* phenotype is reproduced in the *ap1 cal AP1-GR* system (Ler background). Shown are representative buds of *ap1 cal AP1-GR* (top row) *and drmy1 ap1 cal AP1-GR* (bottom row) at day 0 (before DEX induction), day 3 (after 3 DEX inductions, when tissue is collected for RNA, protein, or cytokinin extraction), and day 5 (after 5 DEX inductions). Arrowheads show sepal primordia that are of variable number, position, and sizes. Asterisks indicate periphery of the floral meristem that has limited or no sepal outgrowth. Scale bars, 25 µm.

**(B)** Summary of the inflorescence RNA-seq and proteomics datasets. Shown are numbers of genes in each category. Down, downregulated in *drmy1*; NS, not significantly changed between *drmy1* and WT; Up, upregulated in *drmy1*; NA, not available. Note that in the combined dataset (gene-protein pairs), different genes encoding for the same protein were separately counted, so were different proteins encoded by the same gene. See also Supplemental Dataset 1.

**(C)** Violin and box plots of log2 fold change in RNA between *drmy1* and WT in induced *ap1 cal AP1-GR* inflorescence, for genes encoding ribosomal components (“Structural constituents of the ribosome” GO:0003735, and its offspring terms) and all other genes involved in translation (“Translation” GO:0006412, and its offspring terms). The following genes are labeled on the graph: *UL4Z* (AT3G09630), log2FC = -0.492; *UL4Y* (AT5G02870), log2FC = -0.509; *UL18Z* (AT3G25520), log2FC = -0.459.

**(D-G)** Fluorescence of a constitutively expressed marker supports the hypothesis that *drmy1* has reduced translation rate. (D-F) Representative confocal images of *UBQ10::mCherry-RCI2A* in dissected inflorescences of WT (D), *drmy1* (E), and *ul4y* (F). Numbers show how the signal is divided based on the stage of floral meristem when quantified (IM+1, inflorescence meristem plus stage 1; 2, stage 2; 3, stage 3). Scale bars, 25µm. (G) Signal intensity (i.e. integrated density divided by area) in all images divided as in (D-F). Mean ± SD are shown. Data was fit using a two-way ANOVA model with genotype and stage as two additive factors. Asterisks show statistically significant pairwise contrasts between WT and *drmy1* (p < 2×10^-16^) and between WT and *ul4y* (p = 2.1×10^-15^). Sample sizes: WT IM+1, n = 30; *drmy1* IM+1, n = 22; *ul4y* IM+1, n = 18; WT stage 2, n = 99; *drmy1* stage 2, n = 100; *ul4y* stage 2, n = 52; WT stage 3, n = 39; *drmy1* stage 3, n = 27; *ul4y* stage 3, n = 26.

**(H)** Violin and box plots of log2 fold change in protein level between *drmy1* and WT in induced *ap1 cal AP1-GR* inflorescence, for genes in the same categories as in (C). The following genes are labeled on the graph: *UL4Z* (AT3G09630), log2FC = 0.352; *UL4Y* (AT5G02870), log2FC = 0.811; *UL18Z* (AT3G25520), log2FC = 0.742.

**(I)** Coherent regulation of gene expression by *drmy1* and AZD-8055. Shown is a contingency table of genes downregulated (Down), not significantly changed (NS), and upregulated (Up) in *drmy1* vs WT (columns), and in AZD-8055-treated WT vs mock-treated WT (rows). Bold font shows the number of genes in each category, and gray font shows the expected number of genes if there were no correlation between two conditions (calculated as row margin × column margin / total number of genes). Categories where the number of genes is above expectation are highlighted blue, and categories where the number of genes is below expectation are highlighted red. Chi-square test p < 2.2×10^-16^.

**(J)** Gene ontology enrichment of genes coherently downregulated by both *drmy1* and AZD-8055. Shown are the top 8 terms and their enrichment p-values. Note that the first 7 terms are all related to ribosome and translation. A complete list can be found in Supplementary Dataset 3.

**Supplemental Figure 2. Ribosomal mutations enhance the *drmy1* phenotype, associated with Figure 2.**

**(A-H)** Examples of stage 5 buds from *drmy1* (A), *drmy1 ul4z* (B-D), *drmy1 ul4y* (E-F), and *drmy1 ul18z/+* (G-H). In (B,E,G) sepal primordia within each bud have bigger size differences than typical *drmy1* single mutant buds; asterisks show giant outer sepal primordia and brackets show bud peripheral regions with little or no primordium outgrowth. In (C,F,H), arrowheads show 6 sepal primordia within each bud, which does not occur in *drmy1*. In (D,H), asterisks show the presence of two outer sepal primordia within a bud, instead of one in *drmy1*. Scale bars, 25µm.

**(I-J)** Quantification of sepal primordium number (I) and positional variability (J), comparing each of *drmy1 ul4z* (n = 60), *drmy1 ul4y* (n = 61), and *drmy1 ul18z/+* (n = 69) with *drmy1* (n = 67). “ns” indicates no significant difference in Fisher’s contingency table tests (I) and Wilcoxon’s rank sum tests (J) respectively. Data for *drmy1* is reused from Figure 2H, 2J.

**(K)** Dissected young silique of a *drmy1 ul18z/+* plant. Arrowheads point to aborted ovules. Scale bar, 200 µm.

**Supplemental Figure 3. Sepal primordia in ribosome and TOR mutants catch up in growth to form uniformly sized mature sepals within the bud, associated with Figure 3.**

**(A-F)** Inflorescences (left) of WT (A), *drmy1* (B), *ul4z* (C), *ul4y* (D), *ul18z* (E), and *lst8-1-1* (F), with boxed regions enlarged (right). Blue arrowheads show sepals of regular length, and red arrowheads show sepals shorter than others. Note that sepals in *drmy1* were unable to close due to unequal lengths, while sepals in *ul4z*, *ul4y*, and *ul18z*, and close like in WT. Sepals in *lst8-1-1* were unable to close although there is no apparent variation in length. Scale bars, 0.5 mm.

**(G-L)** Dissected sepals from a bud of WT (G), *drmy1* (H), *ul4z* (I), *ul4y* (J), *ul18z* (K), and two buds of *lst8-1-1* (L). Note that sepals in the *drmy1* bud are of different sizes. Sepals within each bud of *ul4z*, *ul4y*, *ul18z*, and *lst8-1-1* are of similar sizes, although there can be variation between different buds of the same genotype. O, outer sepal. I, inner sepal. L, lateral sepal. Scale bars, 200 µm.

**(M)** Quantification of between-flower variability of sepal length. Length was measured from all imaged sepals of each genotype and each position (outer, inner, lateral), and coefficient of variation (CV) was calculated. A two-sided permutation test (100,000 permutations) for CV difference not equating to zero was done for each pair of genotypes, and results were represented by letters. Sample size: Outer sepal, WT n = 35, *drmy1* n = 43, *ul4z* n = 37, *ul4y* n = 42, *ul18z* n = 39, *lst8-1-1* n = 43. Inner sepal, WT n = 34, *drmy1* n = 46, *ul4z* n = 38, *ul4y* n = 44, *ul18z* n = 37, *lst8-1-1* n = 44. Lateral sepal, WT n = 65, *drmy1* n = 84, *ul4z* n = 81, *ul4y* n = 89, *ul18z* n = 76, *lst8-1-1* n = 82.

**(N)** Quantification of within-flower variability of length. Flowers with length data from at least four sepals were analyzed. A CV of length from all sepals within each flower was calculated, and mean ± SD was plotted, grouped by genotype. A Wilcoxon rank sum test was done for each pair of genotypes, and results were represented by letters. Sample size: WT n = 31 buds, *drmy1* n = 38 buds, *ul4z* n = 33 buds, *ul4y* n = 36 buds, *ul18z* n = 32 buds, *lst8-1-1* n = 39 buds.

**(O-Q)** Live imaging of sepal development from stage 3 to 6 in WT (O), *drmy1* (P), and *ul4y* (Q), showing chlorophyll or propidium iodide channel, and Gaussian curvature of the surface. Note that both *drmy1* and *ul4y* have inner sepals that initiate late (day 2, asterisk). The *drmy1* inner sepal develops slowly, and leaves the bud open at day 3 (red arrowhead). The *ul4y* inner sepal catches up with the rest of the sepals and closes the bud (blue arrowhead). Scale bars, 25 µm.

**Supplemental Figure 4. Inhibition of TOR activity and translation causes auxin maxima formation at variable positions, correlated with variable positions of sepal primordia, associated with Figure 4.**

**(A-E)** Variable patterning of auxin signaling in *drmy1*, *ul4y*, and CHX-treated WT buds corresponds to variable sepal initiation. During the time course, top rows show *DR5::3xVENUS-N7* (yellow), middle rows show composites of *DR5* (yellow) and Chlorophyll (magenta), and bottom rows show Gaussian curvature of the Chlorophyll channel. In the last time point, buds were stained with propidium iodide (top), and Gaussian curvature of the propidium iodide channel is shown on the bottom. (A) In WT, four robustly positioned auxin maxima at day 1 correlates with four robustly positioned sepal primordia at day 4 (blue arrowheads). (B) In *drmy1*, at day 1 there are three robustly positioned auxin maxima (blue arrowheads). At day 2, a diffuse band of auxin signaling occurs in the adaxial periphery of the bud, joining with one of the lateral auxin maxima (red bracket). At day 3, this diffuse band splits into three auxin maxima (red arrowheads), making a total of 5. The maxima correlate with the five sepal primordia at day 4, three at robust positions (blue arrowheads) and two at irregular positions (red arrowheads). (C) In *ul4y*, at day 1 there are two auxin maxima at robust positions (blue arrowheads), one at robust lateral position but much weaker (red arrowhead), and a band of weak auxin signaling in the adaxial periphery of the bud (red bracket). At day 2, the weak auxin maxima at the lateral position got stronger, and the weak band at the adaxial position split into two auxin maxima (red arrowheads). These five auxin maxima correspond to the five sepal primordia at day 3, three in robust positions (blue arrowheads) and two in irregular positions (red arrowheads). (D) In a mock-treated WT bud, four robust auxin maxima at day 6 of the treatment grow into four robust sepal primordia at day 9 (blue arrowheads). (E) In a CHX-treated WT bud, at day 6 there are three stronger auxin maxima (blue arrowheads) and two weaker ones (red arrowheads), corresponding to three bigger regions of outgrowth (blue arrowheads) and two smaller ones (red arrowheads) at day 9. For ease of display, the DR5 channel in CHX-treated WT was brightened three times relative to mock.

**(F-I)** TOR inhibition using Torin2 causes increased cytokinin signaling, and occasional spatial variability in auxin and cytokinin signaling. (F) Late stage 2 buds of WT treated *in vivo* with mock or 2 nmol Torin2 twice a day for 15 days. Shown are *DR5::3xVENUS-N7* in yellow, *TCS::GFP* in cyan, and both merged with propidium iodide in magenta. Note that 3/16 (19%) buds had variable number and position of DR5 and TCS maxima, and 13/16 (81%) had robust DR5 and TCS maxima, although TCS intensity is higher than mock in both cases. (G) Quantification of TCS intensity from maximum intensity projection images, normalized to the mean of WT mock. Shown are mean ± SD. Asterisk shows statistical significance in a two-tailed Student’s t-test (p = 1.2×10^-4^). (H) Circular histograms of DR5 signal distribution (mean ± SD). (I) Circular histograms of TCS signal distribution (mean ± SD). For calculation of circular histograms, please see Figure 4 legends and Materials and Methods. Sample size: WT mock, n = 11 buds; WT Torin2, n = 16 buds. Scale bars in all micrographs, 25 µm.

**Supplemental Figure 5. Translation of uORF-containing ARFs is not universally downregulated in *drmy1*, associated with Figure 5**.

**(A)** *drmy1* has a lower protein-transcript ratio than WT for genes with at least 2 uORFs. 5,086 transcript-protein pairs in our inflorescence dataset were grouped according to the maximum number of uORFs in all transcript isoforms (0, n = 3,485; 1, n = 874; ≥ 2, n = 724) (von Arnim et al., 2014). For each pair, protein-transcript ratio was calculated, log-transformed, and the difference between *drmy1* and WT was plotted. A negative value means this gene has less protein per transcript in *drmy1* than WT, and could indicate reduced translation or protein stability. Medians for each group: 0 uORF, -0.00367; 1 uORF, -0.00808; ≥ 2 uORFs, -0.0243. Asterisk show statistically significant difference from Group 0 in a Wilcoxon rank sum test (p = 3.167×10^-4^), while ns means no significant difference from Group 0 (p = 0.167).

**(B-D)** There is no universal decrease in the expression of uORF-containing ARF reporters. (B) Transcript level of three activator *ARF*s (*ARF5*, *ARF6*, *ARF8*) and two repressor *ARF*s (*ARF3*, *ARF10*) in inflorescence RNA-seq (n = 3 per genotype). *ARF3*, *ARF5*, and *ARF6* contain uORFs before the main ORF, and *ARF8* and *ARF10* do not. Asterisk indicates statistically significant differences between WT and *drmy1* from DESeq2 output. p values: *ARF3*, p = 0.583; *ARF5*, p = 0.497; *ARF6*, p = 0.603; *ARF8*, p = 0.058; *ARF10* p = 0.019. (C) Transcriptional reporters for these ARFs in stage 2 buds of WT and *drmy1* (cyan, GFP or YFP; magenta, propidium iodide). Note that the *pARF3*, *pARF5*, and *pARF6* reporters contain the same uORFs as the genes, reflecting a combination of transcriptional and uORF regulations. Scale bars, 20 µm. (D) Quantification of GFP intensity. Sample size: *pARF3* WT, n = 22; *pARF3 drmy1*, n = 25; *pARF5* WT, n = 22; *pARF5 drmy1*, n = 22; *pARF6* WT, n = 19; *pARF6 drmy1*, n = 28; *pARF8* WT, n = 25; *pARF8 drmy1*, n = 31; *pARF10* WT, n = 20; *pARF10 drmy1*, n = 29. Asterisks show statistically significant differences between WT and *drmy1* in Wilcoxon rank sum tests. p values: *pARF3*, p = 0.3797; *pARF5*, p = 6.22×10^-5^; *pARF6*, p = 2.868×10^-13^; *pARF8*, p = 0.5127; *pARF10* p = 7.073×10^-14^.

**Supplemental Figure 6. Cytokinin signaling causes variability in mature sepal number and size in drmy1, associated with** Figure 5.

Shown are top-view inflorescence images of WT (A), *arr1,10,12* (B), *wol* (C), *drmy1* (D), *drmy1 arr1,10,12* (E), and *drmy1 wol* (F), with boxed areas of individual buds enlarged and shown on the right. In the enlarged views, blue arrowheads point to sepals of regular size, and red arrowheads point to sepals that are much smaller. Scale bars, 0.5 mm.

**Supplemental Figure 7. Investigating other mechanisms that may explain the observed changes in hormone signaling.**

**(A)** Cytokinin abundance does not significantly change in *drmy1*. Shown is mean ± SD of levels of trans-zeatin (tZ), cis-Zeatin (cZ), N^6^-(Δ^2^-Isopentenyl)adenine (iP), trans-Zeatin riboside (tZR), cis-Zeatin riboside (cZR), and N^6^-(Δ^2^-Isopentenyl)adenosine (iPR) quantified by LC-MS in induced WT and *drmy1* inflorescences of *ap1 cal AP1-GR* background. Levels are normalized to the mean tZ level in WT. Sample size: n = 5 for WT; n = 6 for *drmy1*. ns, no significant difference between WT and *drmy1* in two-sided Wilcoxon rank sum tests. P-values: tZ, p = 0.2468; cZ, p = 0.7922; iP, p = 0.2468; tZR, p = 0.1775; cZR, p = 0.6623; iPR, p = 0.6623.

**(B)** Expression of cytokinin signaling components in WT vs *drmy1* inflorescences of *ap1 cal AP1-GR* background (from Supplemental Dataset 1).

**(C-D)** The *ARR7-llama GFP-nes* reporter responds to externally applied cytokinin. Shown are GFP images of the same bud before (C) or after (D) 5 hours of 200 µM BAP treatment. Images are representative of n = 9 buds from two independent lines.

**(E-H)** CHX (E-F) or AZD-8055 (G-H) treatments do not change the subcellular localization of GFP-nes. Images are representative of n = 10 buds (E), n = 9 buds (F), n = 10 buds (G), and n = 11 buds (H). For (C-H), each image was brightened to reveal patterns of GFP distribution. A square region taken from the image containing 5-10 cells is enlarged and shown on the top right. Within the square, GFP intensity was quantified along the dotted line and plotted on the bottom right. X-axis, pixels (range 0-238). Y-axis, GFP intensity (smoothened by taking the average intensity of 11-pixel neighborhoods; range 90-175 in gray value). Scale bars, 25 µm.

**(I-J)** Response of *pAHP3::AHP3-GFP* (I) and *pUBQ10::mCherry-RCI2A* (J) to mock, CHX, and AZD-8055 treatments for 72 hours. Scale bars, 25 µm. For (I), images are representative of n = 13 (mock), n = 15 (CHX), and n = 16 (AZD-8055) buds in two experiments. For (J), images are representative of n = 12 (mock), n = 9 (CHX), and n = 10 (AZD-8055) buds in two experiments.

**(K-N)** The *ARR7-llama GFP-nes* construct partially rescues the mature sepal variability in *drmy1*. Shown are inflorescence images of WT (K), *ARR7-llama GFP-nes* (L), *drmy1* (M), and *drmy1 ARR7-llama GFP-nes* (N). The boxed regions were enlarged and shown on the right of each panel. Note that while *drmy1* buds have normal-sized (blue arrowheads) and smaller (red arrowheads) sepals, some buds in *drmy1 ARR7-llama GFP-nes* have robustly sized sepals (N, middle) while others still show variability (N, right). Scale bars, 0.5 mm.

**(O-P)** *drmy1* has decreased and disrupted pattern of DII degradation. **(O)** Representative images of WT and *drmy1* showing *DII-n3xVENUS* (cyan), *mDII-ntdTomato* (magenta), and merge. For ease of display, the *VENUS* channel was brightened 3 times relative to the *tdTomato* channel. Scale bars, 25 µm. **(P)** Quantification of VENUS/tdTomato ratio. A background of 6 gray value per pixel (determined in blank regions) were subtracted before calculation of ratios. Sample size: WT, n = 8 buds; *drmy1*, n = 19 buds. Asterisk shows statistically significant difference in a Wilcoxon rank sum test (p = 0.01335).

**Supplemental Dataset 1. Inflorescence RNA-seq and proteomics.**

**Supplemental Dataset 2. Unprocessed ribosome profiles.**

**Supplemental Dataset 3. Seedling RNA-seq.**

**Supplemental Dataset 4. Data used in graphs.**

**Supplemental Dataset 5. DNA sequences.**

