## Supplemental Figures for "DRMY1 promotes robust morphogenesis by sustaining the translation of cytokinin signaling inhibitor proteins"

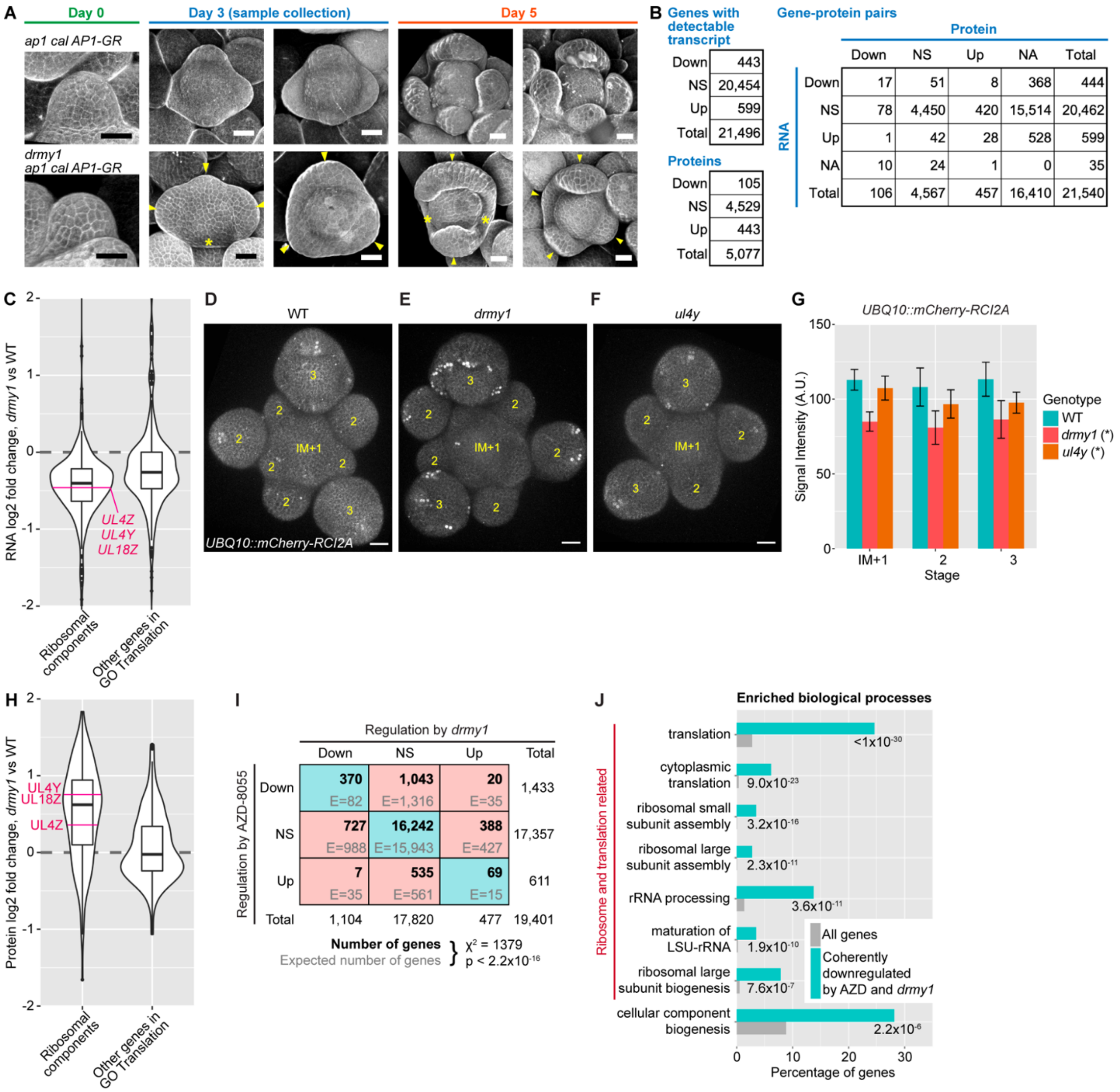


**Supplemental Figure 1. Evidence that the *drmy1* mutant has ribosomal and translation defects, associated with Figure 1.**

**(A)** The *drmy1* phenotype is reproduced in the *ap1 cal AP1-GR* system (Ler background). Shown are representative buds of *ap1 cal AP1-GR* (top row) *and drmy1 ap1 cal AP1-GR* (bottom row) at day 0 (before DEX induction), day 3 (after 3 DEX inductions, when tissue is collected for RNA, protein, or cytokinin extraction), and day 5 (after 5 DEX inductions). Arrowheads show sepal primordia that are of variable number, position, and sizes. Asterisks indicate periphery of the floral meristem that has limited or no sepal outgrowth. Scale bars, 25 µm.

**(B)** Summary of the inflorescence RNA-seq and proteomics datasets. Shown are numbers of genes in each category. Down, downregulated in *drmy1*; NS, not significantly changed between *drmy1* and WT; Up, upregulated in *drmy1*; NA, not available. Note that in the combined dataset (gene-protein pairs), different genes encoding for the same protein were separately counted, so were different proteins encoded by the same gene. See also Supplemental Dataset 1.

**(C)** Violin and box plots of log2 fold change in RNA between *drmy1* and WT in induced *ap1 cal AP1-GR* inflorescence, for genes encoding ribosomal components ("Structural constituents of the ribosome" GO:0003735, and its offspring terms) and all other genes involved in translation ("Translation" GO:0006412, and its offspring terms). The following genes are labeled on the graph: *UL4Z* (AT3G09630), log2FC = -0.492; *UL4Y* (AT5G02870), log2FC = -0.509; *UL18Z* (AT3G25520), log2FC = -0.459.

**(D-G)** Fluorescence of a constitutively expressed marker supports the hypothesis that *drmy1* has reduced translation rate. (D-F) Representative confocal images of *UBQ10::mCherry-RCI2A* in dissected inflorescences of WT (D), *drmy1* (E), and *ul4y* (F). Numbers show how the signal is divided based on the stage of floral meristem when quantified (IM+1, inflorescence meristem plus stage 1; 2, stage 2; 3, stage 3). Scale bars, 25μm. (G) Signal intensity (i.e. integrated density divided by area) in all images divided as in (D-F). Mean $\pm$ SD are shown. Data was fit using a two-way ANOVA model with genotype and stage as two additive factors. Asterisks show statistically significant pairwise contrasts between WT and *drmy1* ($p < 2\times10^{-16}$) and between WT and *ul4y* ($p = 2.1\times10^{-15}$). Sample sizes: WT IM+1, n = 30; *drmy1* IM+1, n = 22; *ul4y* IM+1, n = 18; WT stage 2, n = 99; *drmy1* stage 2, n = 100; *ul4y* stage 2, n = 52; WT stage 3, n = 39; *drmy1* stage 3, n = 27; *ul4y* stage 3, n = 26.

**(H)** Violin and box plots of log2 fold change in protein level between *drmy1* and WT in induced *ap1 cal AP1-GR* inflorescence, for genes in the same categories as in (C). The following genes are labeled on the graph: *UL4Z* (AT3G09630), log2FC = 0.352; *UL4Y* (AT5G02870), log2FC = 0.811; *UL18Z* (AT3G25520), log2FC = 0.742.

**(I)** Coherent regulation of gene expression by *drmy1* and AZD-8055. Shown is a contingency table of genes downregulated (Down), not significantly changed (NS), and upregulated (Up) in *drmy1* vs WT (columns), and in AZD-8055-treated WT vs mock-treated WT (rows). Bold font shows the number of genes in each category, and gray font shows the expected number of genes if there were no correlation between two conditions (calculated as row margin × column margin / total number of genes). Categories where the number of genes is above expectation are highlighted blue, and categories where the number of genes is below expectation are highlighted red. Chi-square test $p < 2.2\times10^{-16}$.

**(J)** Gene ontology enrichment of genes coherently downregulated by both *drmy1* and AZD-8055. Shown are the top 8 terms and their enrichment p-values. Note that the first 7 terms are all related to ribosome and translation. A complete list can be found in Supplementary Dataset 3.

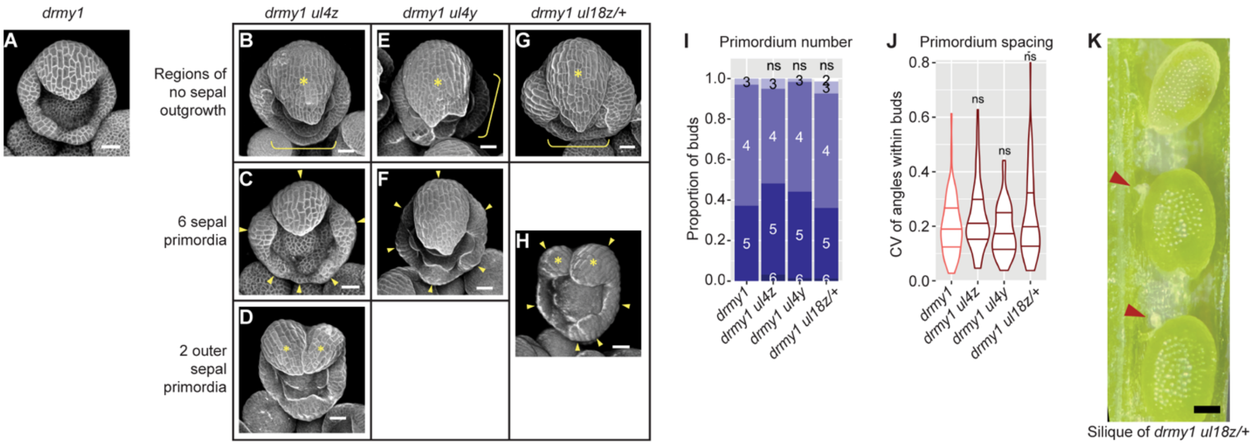


**Supplemental Figure 2. Ribosomal mutations enhance the *drmy1* phenotype, associated with Figure 2.**

**(A-H)** Examples of stage 5 buds from *drmy1* (A), *drmy1 ul4z* (B-D), *drmy1 ul4y* (E-F), and *drmy1 ul18z/+* (G-H). In (B,E,G) sepal primordia within each bud have bigger size differences than typical *drmy1* single mutant buds; asterisks show giant outer sepal primordia and brackets show bud peripheral regions with little or no primordium outgrowth. In (C,F,H), arrowheads show 6 sepal primordia within each bud, which does not occur in *drmy1*. In (D,H), asterisks show the presence of two outer sepal primordia within a bud, instead of one in *drmy1*. Scale bars, 25μm.

**(I-J)** Quantification of sepal primordium number (I) and positional variability (J), comparing each of *drmy1 ul4z* (n = 60), *drmy1 ul4y* (n = 61), and *drmy1 ul18z/+* (n = 69) with *drmy1* (n = 67). “ns” indicates no significant difference in Fisher’s contingency table tests (I) and Wilcoxon’s rank sum tests (J) respectively. Data for *drmy1* is reused from Figure 2H, 2J.

**(K)** Dissected young silique of a *drmy1 ul18z/+* plant. Arrowheads point to aborted ovules. Scale bar, 200 μm.

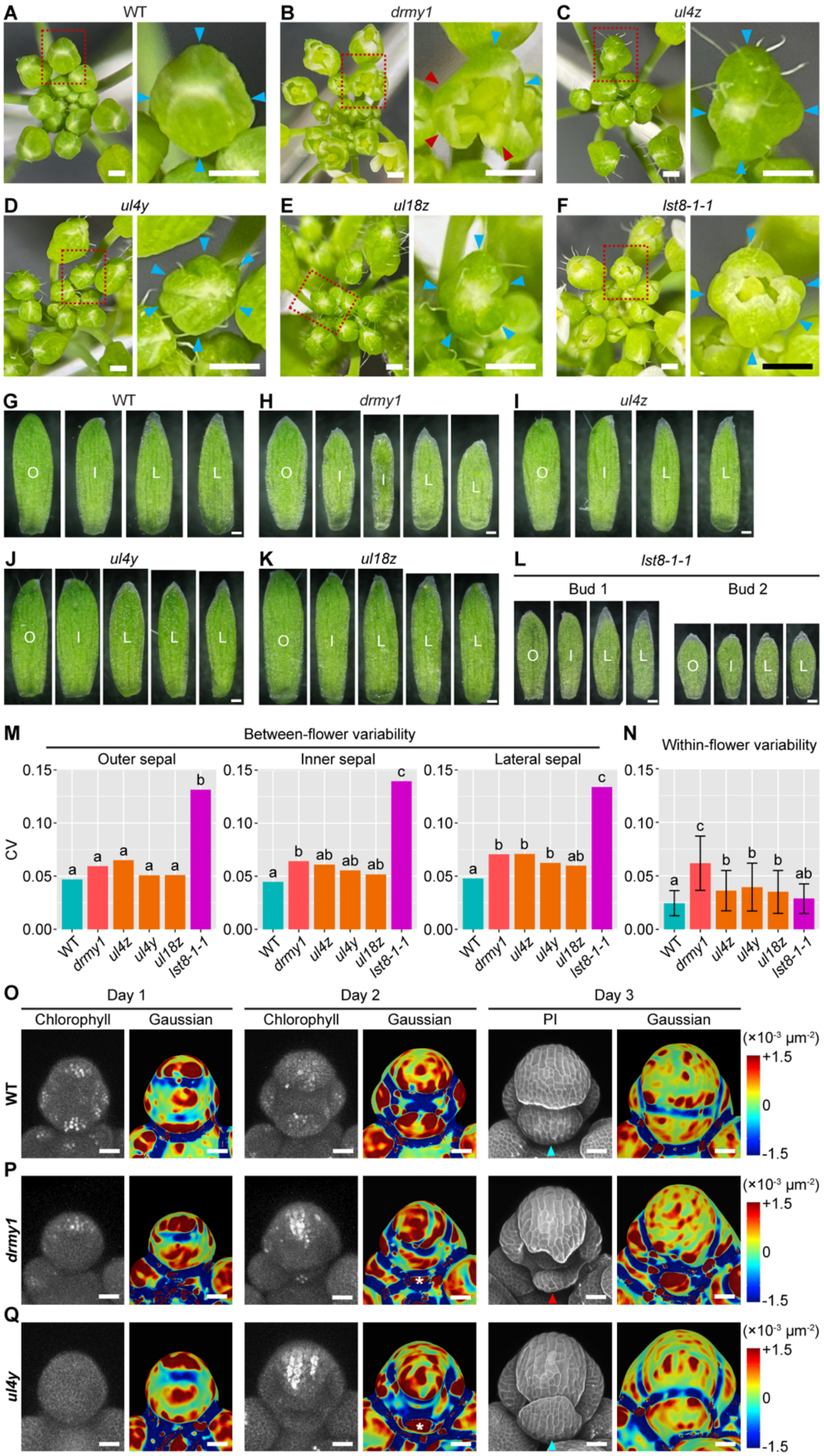


**Supplemental Figure 3. Sepal primordia in ribosome and TOR mutants catch up in growth to form uniformly sized mature sepals within the bud, associated with Figure 3.**

**(A-F)** Inflorescences (left) of WT (A), *drmy1* (B), *ul4z* (C), *ul4y* (D), *ul18z* (E), and *lst8-1-1* (F), with boxed regions enlarged (right). Blue arrowheads show sepals of regular length, and red arrowheads show sepals shorter than others. Note that sepals in *drmy1* were unable to close due to unequal lengths, while sepals in *ul4z*, *ul4y*, and *ul18z*, and close like in WT. Sepals in *lst8-1-1* were unable to close although there is no apparent variation in length. Scale bars, 0.5 mm.
**(G-L)** Dissected sepals from a bud of WT (G), *drmy1* (H), *ul4z* (I), *ul4y* (J), *ul18z* (K), and two buds of *lst8-1-1* (L). Note that sepals in the *drmy1* bud are of different sizes. Sepals within each bud of *ul4z*, *ul4y*, *ul18z*, and *lst8-1-1* are of similar sizes, although there can be variation between different buds of the same genotype. O, outer sepal. I, inner sepal. L, lateral sepal. Scale bars, 200 µm.
**(M)** Quantification of between-flower variability of sepal length. Length was measured from all imaged sepals of each genotype and each position (outer, inner, lateral), and coefficient of variation (CV) was calculated. A two-sided permutation test (100,000 permutations) for CV difference not equating to zero was done for each pair of genotypes, and results were represented by letters. Sample size: Outer sepal, WT n = 35, *drmy1* n = 43, *ul4z* n = 37, *ul4y* n = 42, *ul18z* n = 39, *lst8-1-1* n = 43. Inner sepal, WT n = 34, *drmy1* n = 46, *ul4z* n = 38, *ul4y* n = 44, *ul18z* n = 37, *lst8-1-1* n = 44. Lateral sepal, WT n = 65, *drmy1* n = 84, *ul4z* n = 81, *ul4y* n = 89, *ul18z* n = 76, *lst8-1-1* n = 82.
**(N)** Quantification of within-flower variability of length. Flowers with length data from at least four sepals were analyzed. A CV of length from all sepals within each flower was calculated, and mean ± SD was plotted, grouped by genotype. A Wilcoxon rank sum test was done for each pair of genotypes, and results were represented by letters. Sample size: WT n = 31 buds, *drmy1* n = 38 buds, *ul4z* n = 33 buds, *ul4y* n = 36 buds, *ul18z* n = 32 buds, *lst8-1-1* n = 39 buds.
**(O-Q)** Live imaging of sepal development from stage 3 to 6 in WT (O), *drmy1* (P), and *ul4y* (Q), showing chlorophyll or propidium iodide channel, and Gaussian curvature of the surface. Note that both *drmy1* and *ul4y* have inner sepals that initiate late (day 2, asterisk). The *drmy1* inner sepal develops slowly, and leaves the bud open at day 3 (red arrowhead). The *ul4y* inner sepal catches up with the rest of the sepals and closes the bud (blue arrowhead). Scale bars, 25 µm.

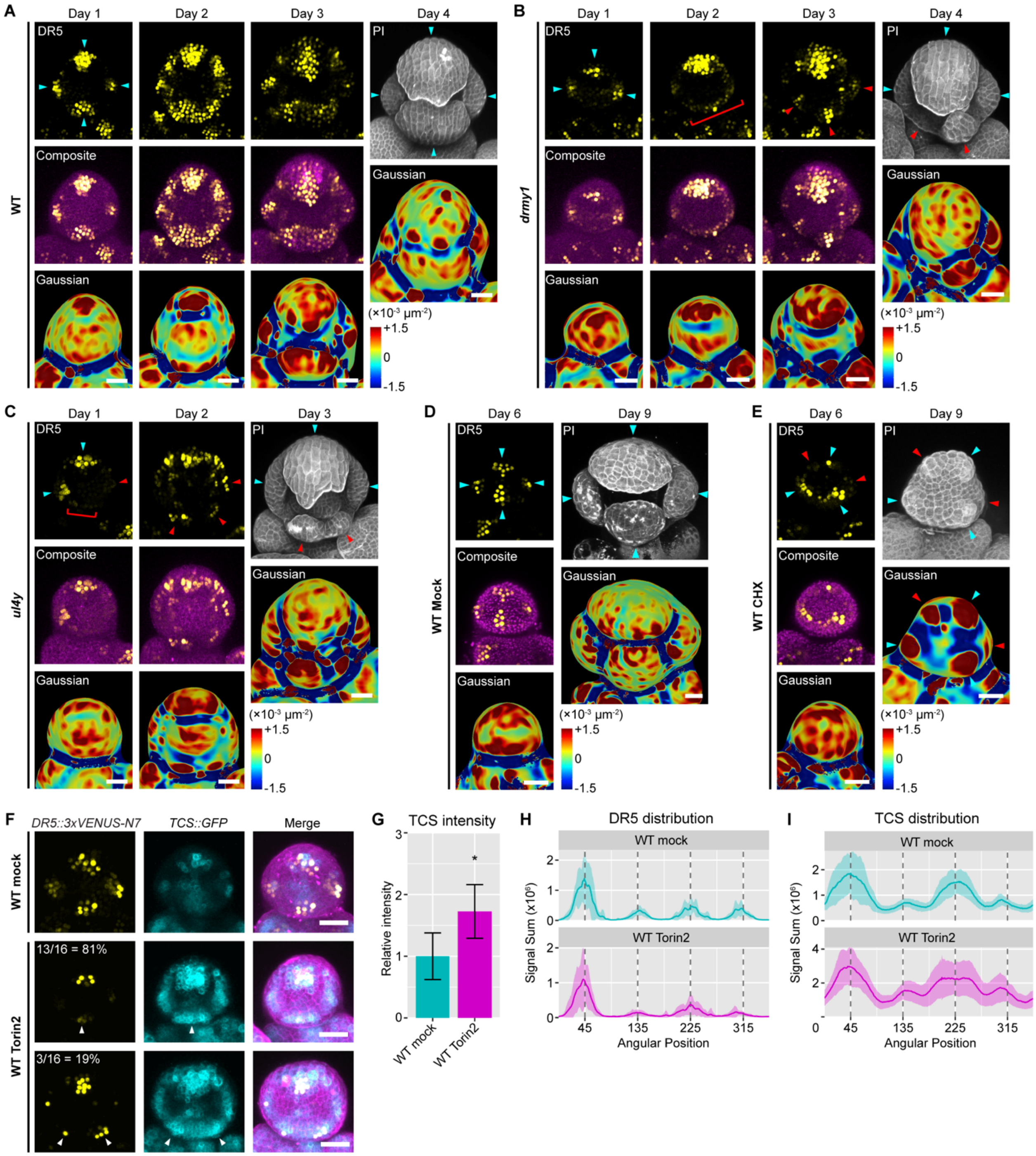


**Supplemental Figure 4. Inhibition of TOR activity and translation causes auxin maxima formation at variable positions, correlated with variable positions of sepal primordia, associated with Figure 4.**

**(A-E)** Variable patterning of auxin signaling in *drmy1*, *ul4y*, and CHX-treated WT buds corresponds to variable sepal initiation. During the time course, top rows show *DR5::3xVENUS-N7* (yellow), middle rows show composites of *DR5* (yellow) and Chlorophyll (magenta), and bottom rows show Gaussian curvature of the Chlorophyll channel. In the last time point, buds were stained with propidium iodide (top), and Gaussian curvature of the propidium iodide channel is shown on the bottom.

(A) In WT, four robustly positioned auxin maxima at day 1 correlates with four robustly positioned sepal primordia at day 4 (blue arrowheads). (B) In *drmy1*, at day 1 there are three robustly positioned auxin maxima (blue arrowheads). At day 2, a diffuse band of auxin signaling occurs in the adaxial periphery of the bud, joining with one of the lateral auxin maxima (red bracket). At day 3, this diffuse band splits into three auxin maxima (red arrowheads), making a total of 5. The maxima correlate with the five sepal primordia at day 4, three at robust positions (blue arrowheads) and two at irregular positions (red arrowheads). (C) In *ul4y*, at day 1 there are two auxin maxima at robust positions (blue arrowheads), one at robust lateral position but much weaker (red arrowhead), and a band of weak auxin signaling in the adaxial periphery of the bud (red bracket). At day 2, the weak auxin maxima at the lateral position got stronger, and the weak band at the adaxial position split into two auxin maxima (red arrowheads). These five auxin maxima correspond to the five sepal primordia at day 3, three in robust positions (blue arrowheads) and two in irregular positions (red arrowheads). (D) In a mock-treated WT bud, four robust auxin maxima at day 6 of the treatment grow into four robust sepal primordia at day 9 (blue arrowheads). (E) In a CHX-treated WT bud, at day 6 there are three stronger auxin maxima (blue arrowheads) and two weaker ones (red arrowheads), corresponding to three bigger regions of outgrowth (blue arrowheads) and two smaller ones (red arrowheads) at day 9. For ease of display, the DR5 channel in CHX-treated WT was brightened three times relative to mock.

**(F-I)** TOR inhibition using Torin2 causes increased cytokinin signaling, and occasional spatial variability in auxin and cytokinin signaling. (F) Late stage 2 buds of WT treated *in vivo* with mock or 2 nmol Torin2 twice a day for 15 days. Shown are *DR5::3xVENUS-N7* in yellow, *TCS::GFP* in cyan, and both merged with propidium iodide in magenta. Note that 3/16 (19%) buds had variable number and position of DR5 and TCS maxima, and 13/16 (81%) had robust DR5 and TCS maxima, although TCS intensity is higher than mock in both cases. (G) Quantification of TCS intensity from maximum intensity projection images, normalized to the mean of WT mock. Shown are mean ± SD. Asterisk shows statistical significance in a two-tailed Student's t-test ($p = 1.2\times10^{-4}$). (H) Circular histograms of DR5 signal distribution (mean ± SD). (I) Circular histograms of TCS signal distribution (mean ± SD). For calculation of circular histograms, please see Figure 4 legends and Materials and Methods. Sample size: WT mock, n = 11 buds; WT Torin2, n = 16 buds. Scale bars in all micrographs, 25 μm.

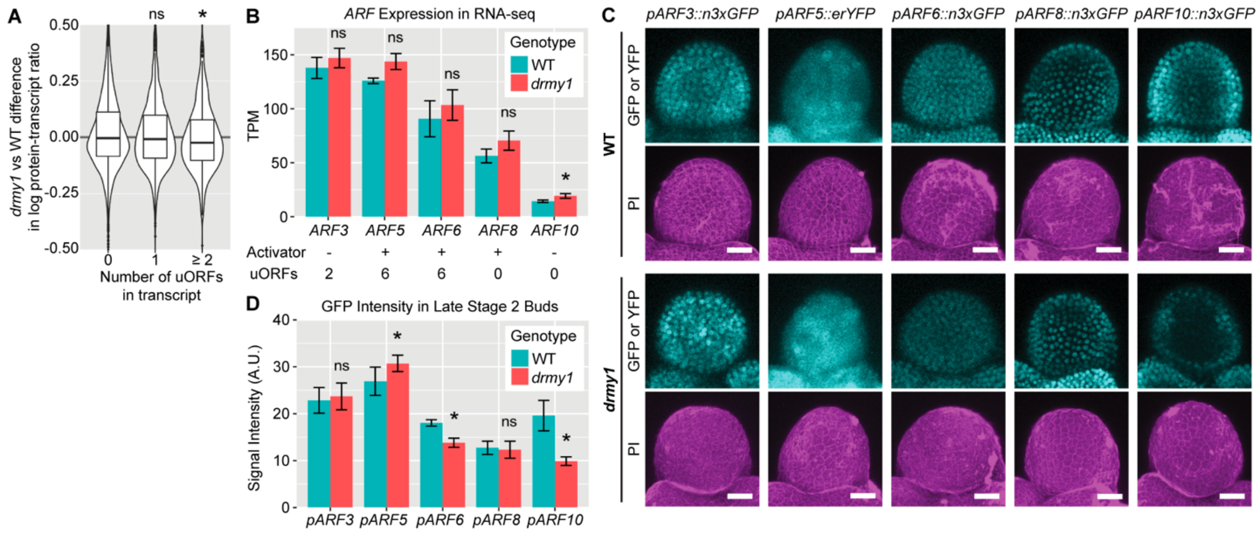


**Supplemental Figure 5. Translation of uORF-containing ARFs is not universally downregulated in *drmy1*, associated with Figure 5.**

**(A)** *drmy1* has a lower protein-transcript ratio than WT for genes with at least 2 uORFs. 5,086 transcript-protein pairs in our inflorescence dataset were grouped according to the maximum number of uORFs in all transcript isoforms (0, n = 3,485; 1, n = 874; ≥ 2, n = 724) (von Arnim et al., 2014). For each pair, protein-transcript ratio was calculated, log-transformed, and the difference between *drmy1* and WT was plotted. A negative value means this gene has less protein per transcript in *drmy1* than WT, and could indicate reduced translation or protein stability. Medians for each group: 0 uORF, -0.00367; 1 uORF, -0.00808; ≥ 2 uORFs, -0.0243. Asterisk show statistically significant difference from Group 0 in a Wilcoxon rank sum test ($p = 3.167\times10^{-4}$), while ns means no significant difference from Group 0 (p = 0.167).

**(B-D)** There is no universal decrease in the expression of uORF-containing ARF reporters. (B) Transcript level of three activator *ARF*s (*ARF5*, *ARF6*, *ARF8*) and two repressor *ARF*s (*ARF3*, *ARF10*) in inflorescence RNA-seq (n = 3 per genotype). *ARF3*, *ARF5*, and *ARF6* contain uORFs before the main ORF, and *ARF8* and *ARF10* do not. Asterisk indicates statistically significant differences between WT and *drmy1* from DESeq2 output. p values: *ARF3*, p = 0.583; *ARF5*, p = 0.497; *ARF6*, p = 0.603; *ARF8*, p = 0.058; *ARF10* p = 0.019. (C) Transcriptional reporters for these ARFs in stage 2 buds of WT and *drmy1* (cyan, GFP or YFP; magenta, propidium iodide). Note that the *pARF3*, *pARF5*, and *pARF6* reporters contain the same uORFs as the genes, reflecting a combination of transcriptional and uORF regulations. Scale bars, 20 µm. (D) Quantification of GFP intensity. Sample size: *pARF3* WT, n = 22; *pARF3 drmy1*, n = 25; *pARF5* WT, n = 22; *pARF5 drmy1*, n = 22; *pARF6* WT, n = 19; *pARF6 drmy1*, n = 28; *pARF8* WT, n = 25; *pARF8 drmy1*, n = 31; *pARF10* WT, n = 20; *pARF10 drmy1*, n = 29. Asterisks show statistically significant differences between WT and *drmy1* in Wilcoxon rank sum tests. p values: *pARF3*, p = 0.3797; *pARF5*, $p = 6.22\times10^{-5}$; *pARF6*, $p = 2.868\times10^{-13}$; *pARF8*, p = 0.5127; *pARF10* $p = 7.073\times10^{-14}$.

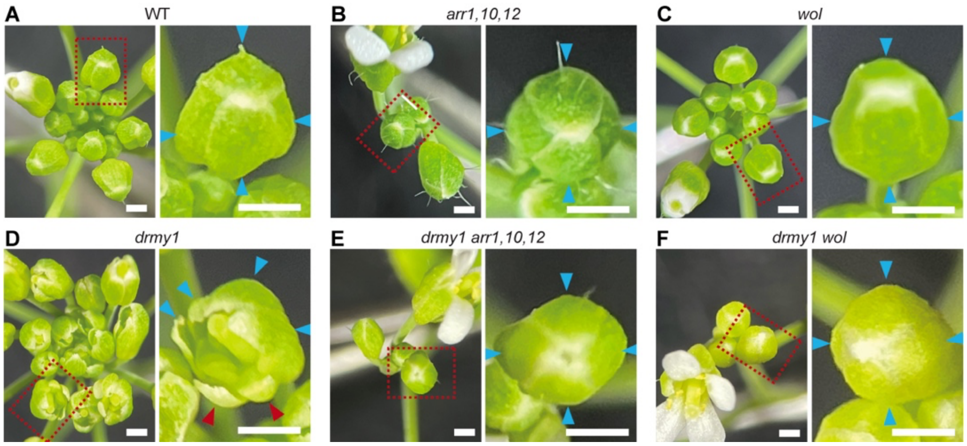


**Supplemental Figure 6. Cytokinin signaling causes variability in mature sepal number and size in *drmy1*, associated with Figure 5.**

Shown are top-view inflorescence images of WT (A), *arr1,10,12* (B), *wol* (C), *drmy1* (D), *drmy1 arr1,10,12* (E), and *drmy1 wol* (F), with boxed areas of individual buds enlarged and shown on the right. In the enlarged views, blue arrowheads point to sepals of regular size, and red arrowheads point to sepals that are much smaller. Scale bars, 0.5 mm.

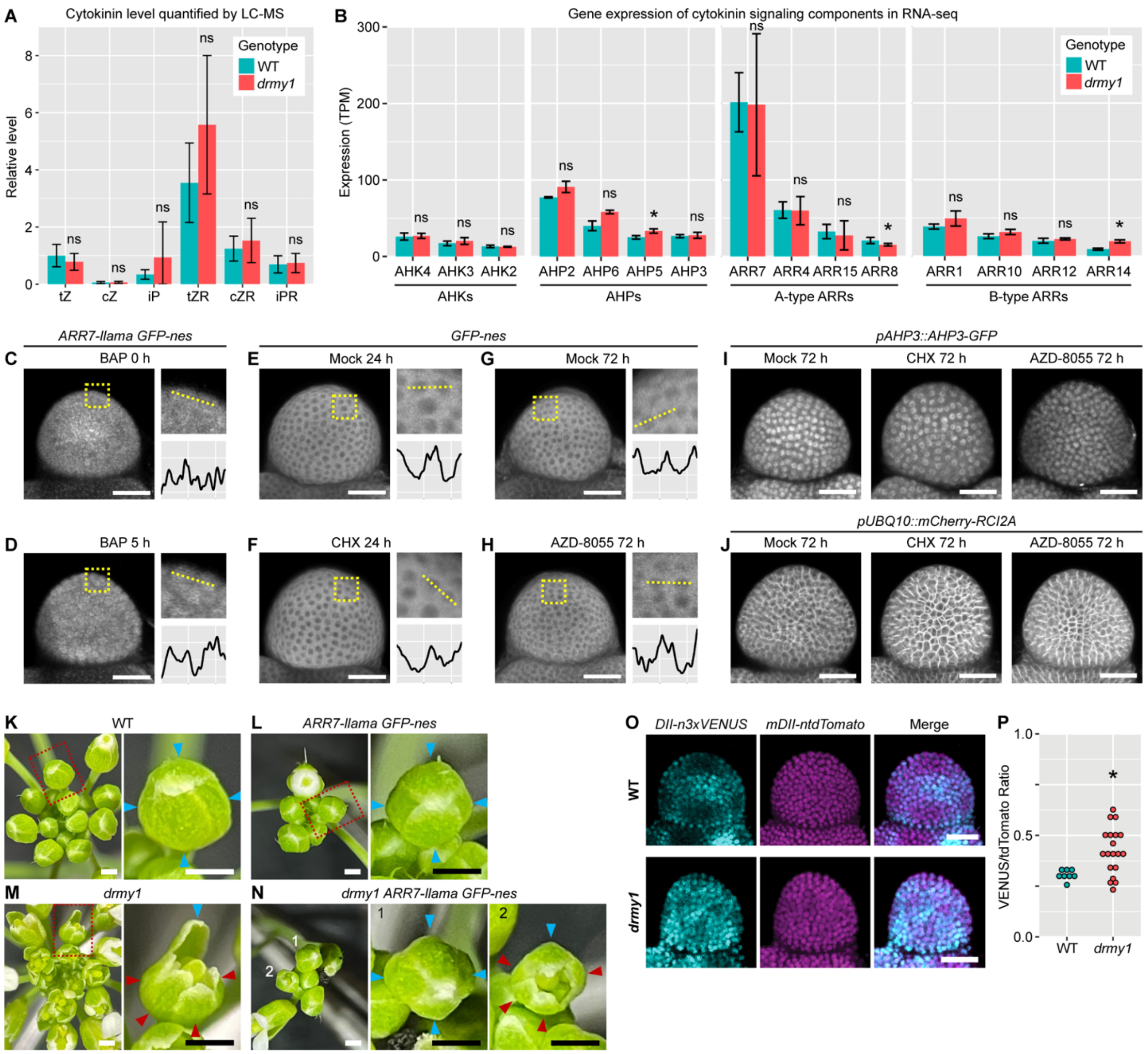


**Supplemental Figure 7. Investigating other mechanisms that may explain the observed changes in hormone signaling.**

**(A)** Cytokinin abundance does not significantly change in *drmy1*. Shown is mean ± SD of levels of trans-zeatin (tZ), cis-Zeatin (cZ), $N^6$-($\Delta^2$-Isopentenyl)adenine (iP), trans-Zeatin riboside (tZR), cis-Zeatin riboside (cZR), and $N^6$-($\Delta^2$-Isopentenyl)adenosine (iPR) quantified by LC-MS in induced WT and *drmy1* inflorescences of *ap1 cal AP1-GR* background. Levels are normalized to the mean tZ level in WT. Sample size: n = 5 for WT; n = 6 for *drmy1*. ns, no significant difference between WT and *drmy1* in two-sided Wilcoxon rank sum tests. P-values: tZ, p = 0.2468; cZ, p = 0.7922; iP, p = 0.2468; tZR, p = 0.1775; cZR, p = 0.6623; iPR, p = 0.6623.

**(B)** Expression of cytokinin signaling components in WT vs *drmy1* inflorescences of *ap1 cal AP1-GR* background (from Supplemental Dataset 1).

**(C-D)** The *ARR7-llama GFP-nes* reporter responds to externally applied cytokinin. Shown are GFP images of the same bud before (C) or after (D) 5 hours of 200 µM BAP treatment. Images are representative of n = 9 buds from two independent lines.
**(E-H)** CHX (E-F) or AZD-8055 (G-H) treatments do not change the subcellular localization of GFP-nes. Images are representative of n = 10 buds (E), n = 9 buds (F), n = 10 buds (G), and n = 11 buds (H). For (C-H), each image was brightened to reveal patterns of GFP distribution. A square region taken from the image containing 5-10 cells is enlarged and shown on the top right. Within the square, GFP intensity was quantified along the dotted line and plotted on the bottom right. X-axis, pixels (range 0-238). Y-axis, GFP intensity (smoothened by taking the average intensity of 11-pixel neighborhoods; range 90-175 in gray value). Scale bars, 25 µm.
**(I-J)** Response of *pAHP3::AHP3-GFP* (I) and *pUBQ10::mCherry-RCI2A* (J) to mock, CHX, and AZD-8055 treatments for 72 hours. Scale bars, 25 µm. For (I), images are representative of n = 13 (mock), n = 15 (CHX), and n = 16 (AZD-8055) buds in two experiments. For (J), images are representative of n = 12 (mock), n = 9 (CHX), and n = 10 (AZD-8055) buds in two experiments.
**(K-N)** The *ARR7-llama GFP-nes* construct partially rescues the mature sepal variability in *drmy1*. Shown are inflorescence images of WT (K), *ARR7-llama GFP-nes* (L), *drmy1* (M), and *drmy1 ARR7-llama GFP-nes* (N). The boxed regions were enlarged and shown on the right of each panel. Note that while *drmy1* buds have normal-sized (blue arrowheads) and smaller (red arrowheads) sepals, some buds in *drmy1 ARR7-llama GFP-nes* have robustly sized sepals (N, middle) while others still show variability (N, right). Scale bars, 0.5 mm.
**(O-P)** *drmy1* has decreased and disrupted pattern of DII degradation. **(O)** Representative images of WT and *drmy1* showing *DII-n3xVENUS* (cyan), *mDII-ntdTomato* (magenta), and merge. For ease of display, the *VENUS* channel was brightened 3 times relative to the *tdTomato* channel. Scale bars, 25 µm. **(P)** Quantification of VENUS/tdTomato ratio. A background of 6 gray value per pixel (determined in blank regions) were subtracted before calculation of ratios. Sample size: WT, n = 8 buds; *drmy1*, n = 19 buds. Asterisk shows statistically significant difference in a Wilcoxon rank sum test (p = 0.01335).
