## Supplementary figures and images for "DRMY1 promotes robust morphogenesis by sustaining the translation of cytokinin signaling inhibitor proteins"

### Suppl Dataset 2 Unprocessed polysome profiles.pdf

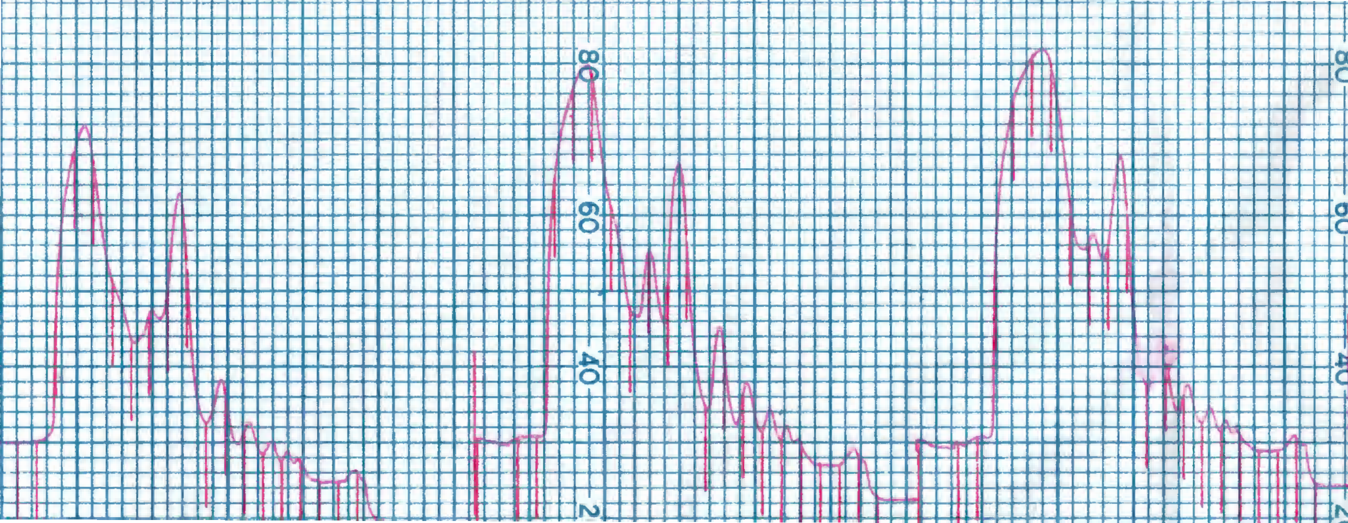

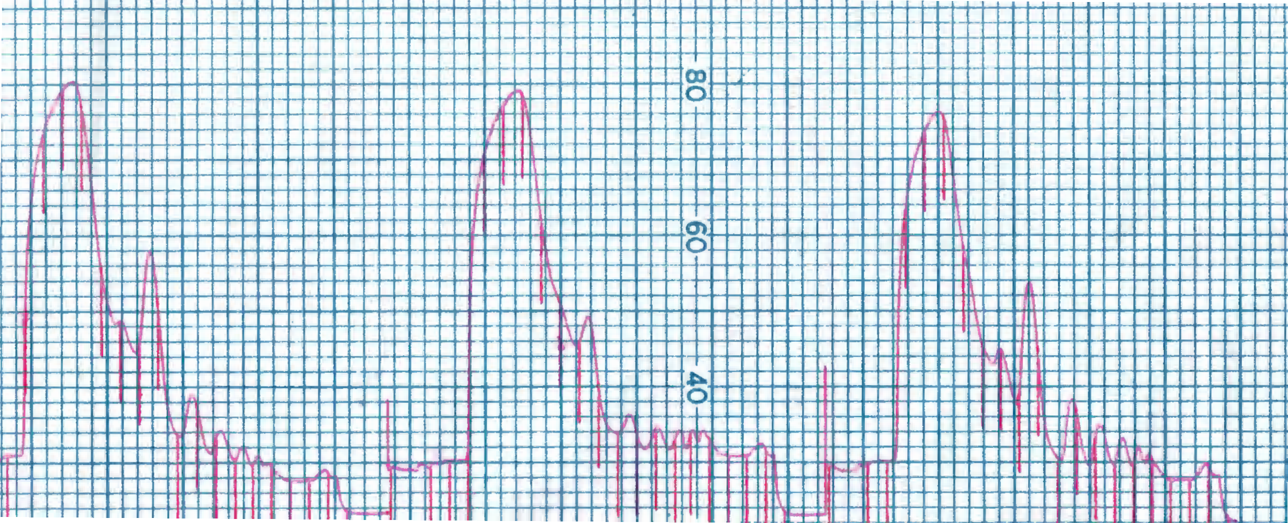
